# SBGN Bricks Ontology as a tool to describe recurring concepts in molecular networks

**DOI:** 10.1101/2020.11.16.369330

**Authors:** Adrien Rougny, Vasundra Touré, John Albanese, Dagmar Waltemath, Denis Shirshov, Anatoly Sorokin, Gary D. Bader, Michael L. Blinov, Alexander Mazein

## Abstract

A comprehensible representation of a molecular network is key to communicating and understanding scientific results in systems biology. The Systems Biology Graphical Notation (SBGN) has emerged as the main standard to represent such networks graphically. It has been implemented by different software tools, and is now largely used to communicate maps in scientific publications. However, learning the standard, and using it to build large maps, can be tedious. Moreover, SBGN maps are not grounded on a formal semantic layer and therefore do not enable formal analysis. Here, we introduce a new set of patterns representing recurring concepts encountered in molecular networks, called SBGN bricks. The bricks are structured in a new ontology, the BricKs Ontology (BKO), to define clear semantics for each of the biological concepts they represent. We show the usefulness of the bricks and BKO for both the template-based construction and the semantic annotation of molecular networks. The SBGN bricks and BKO can be freely explored and downloaded at sbgnbricks.org.

## 1 Introduction

To better understand how complex biological systems work, we need to represent our knowledge in a clear and unambiguous form that is accessible to both scientists and computational agents. These representations form the basis for mathematical modelling and provides a prior-knowledge view for high-throughput data analysis, interpretation and hypothesis generation [16, 30]. The Systems Biology Graphical Notation (SBGN) [21] was developed as a standard for the graphical representation of molecular networks. It is composed of three complementary languages: Process Description (PD) [26] represents modulated processes such as catalyzed reactions; Activity Flow (AF) [23] represents biomolecular activities and the influences they have on each other; and Entity Relationship (ER) [31] represents relationships between biomolecular entities such as molecular interactions. Each language introduces a fixed set of glyphs (i.e. standardized symbols) that represent well-defined biological or bio-molecular elementary concepts (e.g. a macromolecule, a stoichiometric process, a stimulation). Such glyphs can be assembled to form complex SBGN diagrams. In these diagrams, one can also identify representations of less elementary concepts that are recurrent in molecular networks, such as complex formations, reversible metabolic reactions, or protein phosphorylations. One diagram may contain several specific occurrences of such concepts. For example, a signalling pathway typically exhibits multiple protein phosphorylations constituting a signalling cascade, and a metabolic pathway exhibits several chained metabolic reactions. Similar higher order composite structures, often called templates,idioms or patterns appear in other formal modelling languages like circuit diagrams or UML as well as computer programming languages.

In the first SBGN bricks paper [15], we showed that these occurrences could be generalised into generic templates called SBGN bricks, which could then be used as building blocks when building SBGN diagrams. We defined several of such templates in the three SBGN languages and showed how the building blocks were applied for educational purposes. Furthermore, SBGN bricks were used to speed-up the creation of SBGN diagrams via template-based construction [15]. The template-based construction was implemented for SBGN-ED [8] and PathVisio [35, 18] (sbgnbricks.sourceforge.net) and is also supported by more recent editors such as Newt [27] (newteditor.org) and Krayon (github.com/draeger-lab/krayon4sbgn).

In this manuscript, we introduce a comprehensive set of SBGN bricks that largely extends the ones previously introduced in [15], as well as a novel approach to defining a semantic layer which structures the set of SBGN bricks. We first refine the concept of SBGN bricks by making a new distinction between template bricks and instance bricks. We define a template brick as a graphical pattern representing a certain biomolecular process or activity (e.g. a protein phosphorylation, a protein kinase activity). Template bricks may, for example, be used to generate or match instance bricks which represent specific instances of the bio-molecular process or activity defined in the template brick itself (e.g. the phosphorylation of the RSK protein). We then introduce a new ontology, the BricK Ontology (BKO), that structures the set of template bricks by associating them to well-defined biomolecular concepts imported from the Gene Ontology [1, 5] and the Systems Biology Ontology [6]. We show how the ontology is implemented and how it can be navigated from a dedicated website (sbgnbricks.org). Finally, we evaluate the completeness of our new set of template bricks with an in-depth analysis of the nature of all instance bricks matched by this set in the maps of the Atlas of Cancer Signalling Network [17] and the PANTHER database [32, 22].

## 2 Results

In Figure 1 we used SBGN bricks to identify recurring concepts in biological networks. Figure 1 shows an SBGN PD map representing the Insulin/IGF pathway (adapted from identifiers.org/panther.pathway:P00032), annotated by a number of terms that describe generic concepts, such as “protein phosphorylation”. Each coloured box represent an instance brick and bricks cover the whole pathway. They are instantiated from a reduced number of more generic patterns, that we call template bricks. For example, the three green bricks represent specific occurrences of a “protein phosphorylation” generic pattern, or template brick, depicting “a process glyph linked to an unphosphorylated macromolecule glyph via a consumption arc, and to a phosphorylated macromolecule glyph carrying the same label via a production arc”. Hence, one can identify bricks that are instances of more generic template bricks, which in turn may be associated with terms describing generic concepts. In this sense, template bricks may be viewed as canonical representations of generic concepts. Continuing with our example, the green bricks constituting the map of Figure 1 are therefore instances of a template brick that is a canonical representation of the concept described by the term “protein phosphorylation”. Two different template bricks may also be associated with the same term, because a concept may have two or more canonical representations, depending on the context. For example, in the map on Figure 1, the orange bricks and the dark blue brick are all specific occurrences of a “stimulatory activity”. However, the orange bricks describe a stimulation on a process glyph while the dark blue brick describes a stimulation on a phenotype glyph, meaning that they are instances of two different template bricks.

**Figure 1:**
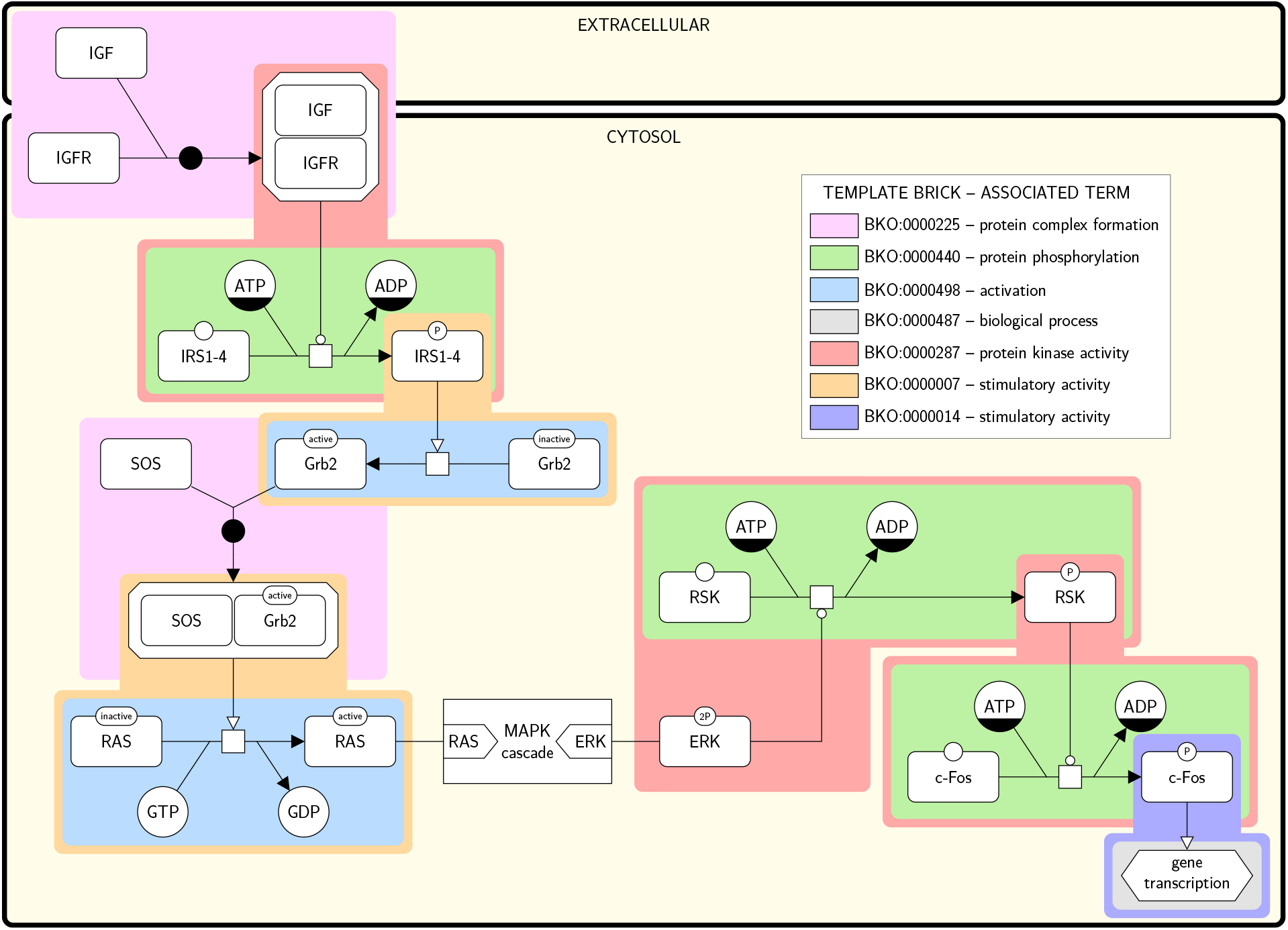
SBGN PD map of the Insulin/IGF pathway-mitogen activated protein kinase/MAP kinase cascade annotated with terms using template bricks. The map is adapted from identifiers.org/panther.pathway:P00032 and uses a submap to hide the MAPK cascade. Colored boxes surround individual instance bricks matched by the template bricks given in the legend. The color of the surrounding box identifies the template brick the instance is matched by (e.g., the template brick in green (BKO:0000440) representing a protein phosphorylation matches three different instances in the maps). Each template brick is associated with a term that describes a biological concept (biomolecular process or activity). Two different template bricks may be associated to the same term, e.g. the template brick in orange (BKO:0000007) and the one in purple (BKO:0000014) are both associated to the term “stimulatory activity”, the first one representing the stimulation of a stoichiometric process, and the second one the stimulation of a phenotype.

We built a comprehensive set of 476 template bricks that substantially extends the one introduced by Junker *et al*. in 2012 [15]. To better structure this set and to precisely describe the complex relationships the bricks share with the concepts they represent, we coupled them with a new ontology, the BricKs Ontology (BKO), that gathers and organizes all bricks and terms. We then evaluated the completeness of our set of template bricks by annotating all SBGN PD maps of the ACSN [17] and PANTHER [32, 22] databases.

### 2.1 Template bricks

A template brick is a graphical pattern representing a biological concept described by a term. It may be used for both generating particular SBGN representations (template-based construction) and searching for such representations in actual SBGN maps (annotation). A particular SBGN representation generated from or matched by a template brick is called an instance brick, and we say that it is an instance of that template brick. Figure 2 gives an example of a template brick and two of its possible instances, taken from the map of Figure 1 (in pink). The template brick is a PD representation of the concept described by the term “protein kinase activity,” whereas the two instance bricks are two PD representations of specific occurrences of this concept: the first one represents the kinase activity of ERK (which catalyzes the phosphorylation of RSK), while the second one represents the kinase activity of the complex IGF/IGFR (which catalyzes the phosphorylation of IRS1-4). Each of these instance bricks also contains an instance brick of another template brick representing the term “protein phosphorylation” (in green).

**Figure 2:**
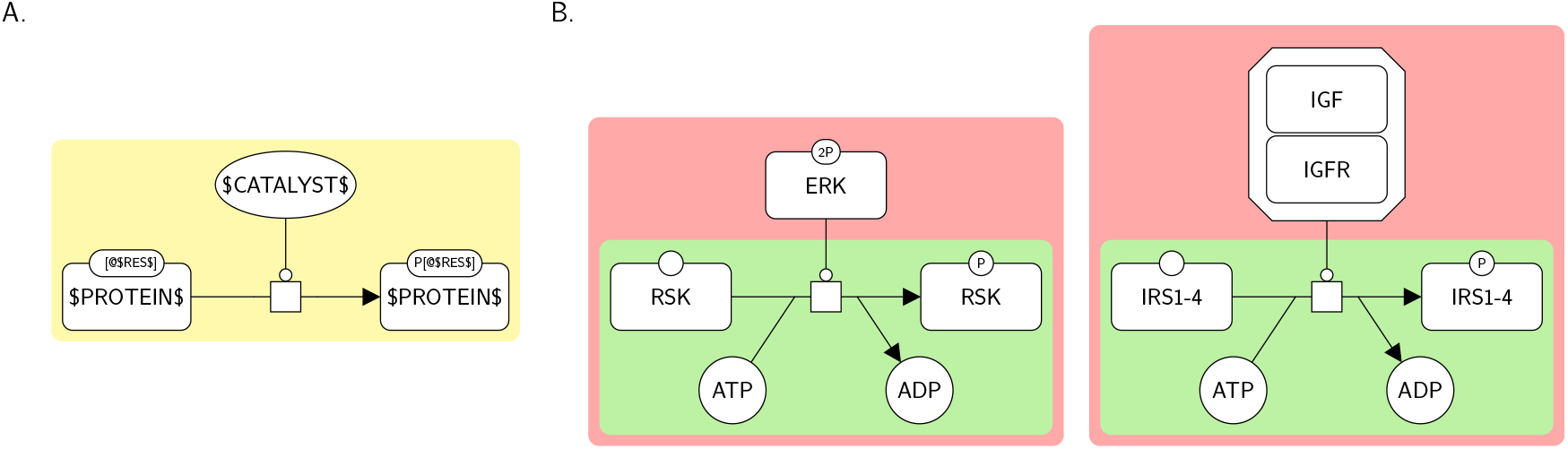
A template brick and two examples of its instance bricks. **A**. A template brick (BKO:0000287) representing the term “protein kinase activity” (GO:0004672). $LABEL$ can represent, or match, any string. **B**. Two instances of this template brick (in pink), taken from Figure 1. Each of these instance bricks also contains an instance brick of another template brick representing the term “protein phosphorylation” (in green).

Template bricks are described using the SBGN languages (here in Figure 2, PD) and additional textual constructs contained in labels that allow, for instance, expressing generality or repetition (see the Methods section for more details).

### 2.2 Classification of bricks in the BricK Ontology

We built an ontology to formally structure the set of bricks, the BricKs Ontology (BKO). Figure 3 displays an excerpt of BKO, showing two template bricks associated with the term “modulatory activity” and that are used in Figure 1. BKO allows organizing the set of template bricks and associating them formally to well-defined concepts (both imported from GO and SBO, and newly defined). It opens a whole set of new possibilities for automated analysis of biological maps, including ontological reasoning. BKO is composed of three main types of elements: terms, template bricks, and categories. The ontology includes template bricks built with the three SBGN languages (PD, AF and ER). In the following, we focus on SBGN PD; specific features of the other two languages will be described later on.

**Figure 3:**
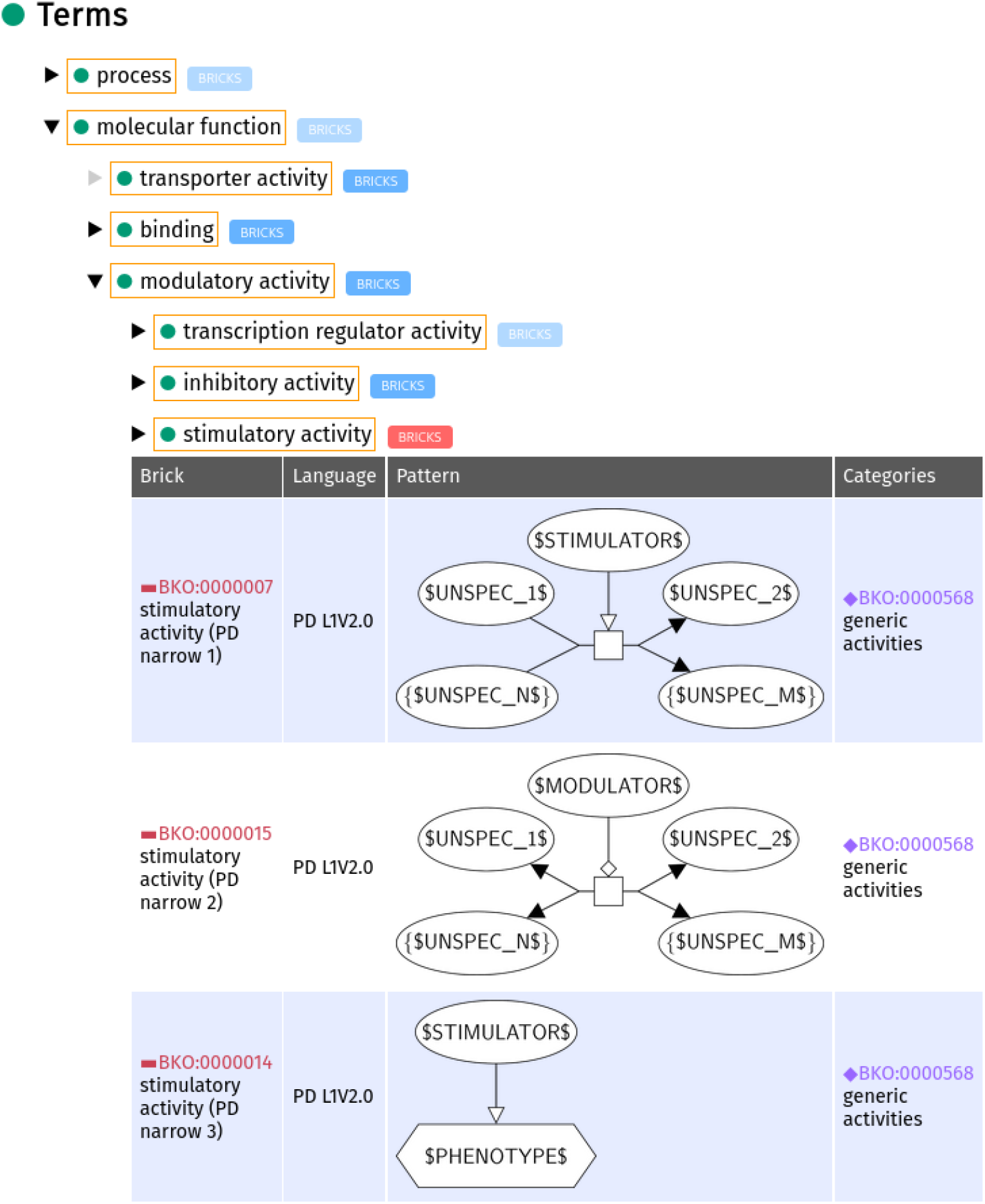
Excerpt of BKO showing three template bricks associated with the term “stimulatory activity”. This excerpt is captured from the SBGN bricks website where the ontology can be navigated (sbgnbricks.org). The parent term “modulatory activity” (BKO:0000059) and the descendant term “stimulatory activity” (BKO:0000075) both have related template bricks associated with them. The presence of these bricks is indicated by a bright blue “BRICKS” label next to the term. When the bricks are displayed, as in the case of “stimulatory activity”, the box containing “BRICKS” turns to a red colour. Some terms, such as “molecular function” (GO:0003674) do not have any associated PD bricks, denoted by the pale blue “BRICKS” label. The pale grey arrow to the left of the term “transporter activity” (GO:0005215) indicates that the term has no further descendants. Figure 1 includes instances of the first and last template bricks (BKO:0000007 and [BKO:00000014](http://sbgnbricks.org/BKO/BKO:0000001), respectively).

#### 2.2.1 Terms

Each biological or biochemical concept is identified by a term that constitutes a class of the ontology (in the ontology and in the following, we combine terms and concepts). Most of the terms and the ontological relationships they share are adapted from SBO (for “processes”) and GO (for “molecular functions”). Each term without a descendant in the ontology is associated with at least one template brick. Some terms are not associated with any template brick because they cannot be represented using generic patterns in either of the three SBGN languages. Such terms will, however, have a descendant associated with at least one brick, hence their presence in the ontology.

#### 2.2.2 Template bricks

A template brick, formally is a graph that is a valid SBGN diagram under one of its three languages and all of its entity nodes are is a graphical pattern representing a term (see Figure 3 for an example). With regards to the ontology, it may be viewed as an ontological class defined as the set of all its instances. Hence, we built an ontological class for each template brick. We also built all subsumption relations (superclass/subclass relations) they share: a template brick A is a subclass of template brick B if and only if all instances that may be matched by A may also be matched by B.

Each template brick is associated with the term(s) of the ontology it represents, as described further below.

#### 2.2.3 Categories

We offer broad categories to provide an easier way to browse the template bricks. In the ontology, each category constitutes an individual of a generic “category” class and is associated with one or more template bricks using a symmetrical relation. For example, the map on Figure 1 contains multiple instances of the template bricks associated with the terms “protein phosphorylation” (BKO:0000438) and “protein kinase activity” (GO:0004672). Both these template bricks belong to the same category “phosphorylation and dephosphorylation” (BKO:0000574).

#### 2.2.4 Association of template bricks with terms

In the ontology, each template brick is associated with the term it represents using a specific relation.

If a template brick fully represents the concept described by the term, then it is associated with the term using a relation labeled main. In Figure 4 panel A, the template brick fully represents the term “translation” (SBO:0000184). Hence it is associated with this term using the main relation. If, on the other hand, a template brick represents only part of the concept described by the term, and it represents fully only a subconcept that is not described by any term of the ontology, then it is associated to the term with a relation labeled *narrow* (analogously to the *narrow* synonyms of the GO ontology). Usually, this case arises when a concept cannot be represented by a unique template brick, and we do not wish to include in the ontology terms for the subconcepts that could each be represented by a unique template brick because they are not completely relevant in terms of biology, or it would complexify the ontology too much. Figure 4 panel B shows two template bricks associated with the term “stimulatory activity” (BKO:0000003) using the *narrow* relation. The top template brick represents the stimulation of an irreversible stoichiometric process, while the bottom one represents the stimulation of a phenotype. Another example of usage of the *narrow* relation is related to the reversibility of processes. The term “catalytic activity” (GO:0003824) cannot be represented using a unique template brick, because only reversible or irreversible processes can be represented, and there is no way to represent processes that might be indistinctly one or the other.

**Figure 4:**
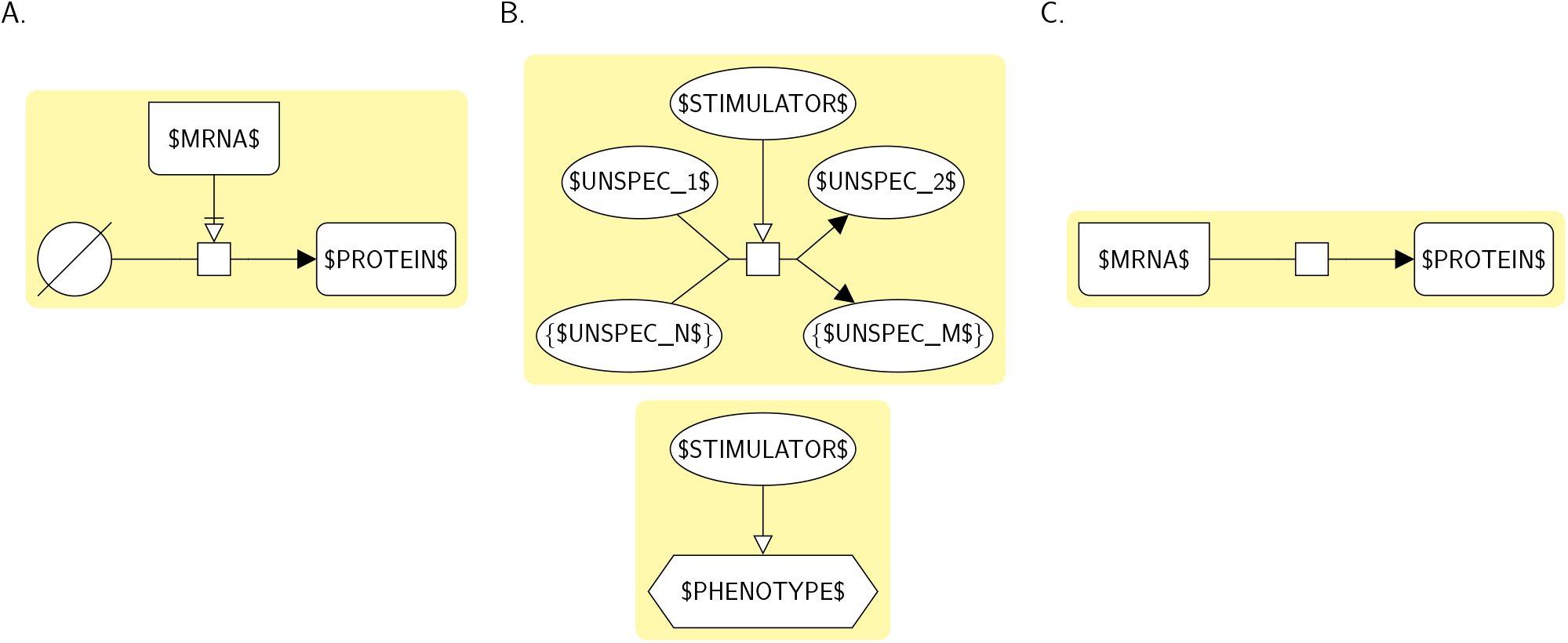
Association of template bricks with terms using the main, *narrow* and *alternate* relations. **A**. A template brick representing the term “translation” (SBO:0000184). It fully represents this term, and is associated with it using the main relation. **B**. Two template bricks representing the term “stimulatory activity” (BKO:0000003). The top brick represents the stimulation of an irreversible process, while the bottom one represents the stimulation of a phenotype. Since each brick only represents a subconcept of the concept described by the term, it is associated with the term using the *narrow* relation. **C**. Another template brick representing the term “translation”. It fully represents the term, but there is another template brick that also fully represents the term and that is more appropriate (see A). Hence it is associated with this term with the *alternate* relation. This template brick was only added to the set for compatibility with translation processes of CellDesigner [11]. Since this representation is not fundamentally correct with respect to the semantics of SBGN PD, it should not be used in template-based construction, but it is relevant for annotation of maps, for example.

However, to avoid having a term for each of the two types of processes, we built two bricks, one for each type of process (BKO:0000004 and BKO:0000005). Each of the bricks represents only part of the term “catalytic activity”, and both bricks are associated with the term using the *narrow* relation.

Finally, if the template brick fully represents the concept described by the term, but there is another template brick also fully representing this concept and that is more relevant or preferred, then the former is associated with the term using a relation labeled *alternate* (while the latter is associated with it using the relation main described above). In Figure 4 panel C, the template brick fully represents the term “translation”. However, the brick of Figure 4 panel A also fully represents this term and is more accurate. Hence, the brick of panel C is associated with the term using the *alternate* relation.

### 2.3 Extending to the ER and AF languages

The PD language allows depicting biochemical processes under a mechanistic and temporal point of view. It is widely used by pathway databases and modelling tools to represent molecular networks under the form of detailed maps [34]. Hence we primarily focused on building a complete set of template bricks expressed in PD. However we also built template bricks expressed in the other two SBGN languages, AF and ER. AF permits representing a flow of activities that influence each other, ER is used to represent relationships between biochemical entities without any temporal aspect. Although the three SBGN languages are orthogonal, some molecular networks may be depicted using one or the other, depending on the point of view one wants to adopt. Figure 5 and Figure 6 give excerpts of the AF and ER maps matching the PD map of Figure 1 and annotated with terms using AF and ER template bricks, respectively. The complete annotated maps are given in Supplementary Figure S1 (AF) and Supplementary Figure S2 (ER). These maps illustrate how template bricks expressed in the three different languages may be aligned. For example, the term “protein kinase activity” (GO:0004672) can be represented in PD, AF and ER, and hence is associated with at least one template brick per language (in pink in Figure 1, Figure 5 and Figure 6). Some terms, however, cannot be represented in all three languages. For example, it is not possible to represent biomolecular processes in AF, and therefore the term “protein phosphorylation” (BKO:0000438) is only associated with PD and ER template bricks. By associating template bricks with terms, BKO aligns representations expressed in the three SBGN languages, based on the concepts they represent. This alignment is further illustrated in Supplementary Table S1, showing the PD, AF and ER template bricks associated with the terms describing some of the most common concepts encountered in molecular networks.

**Figure 5:**
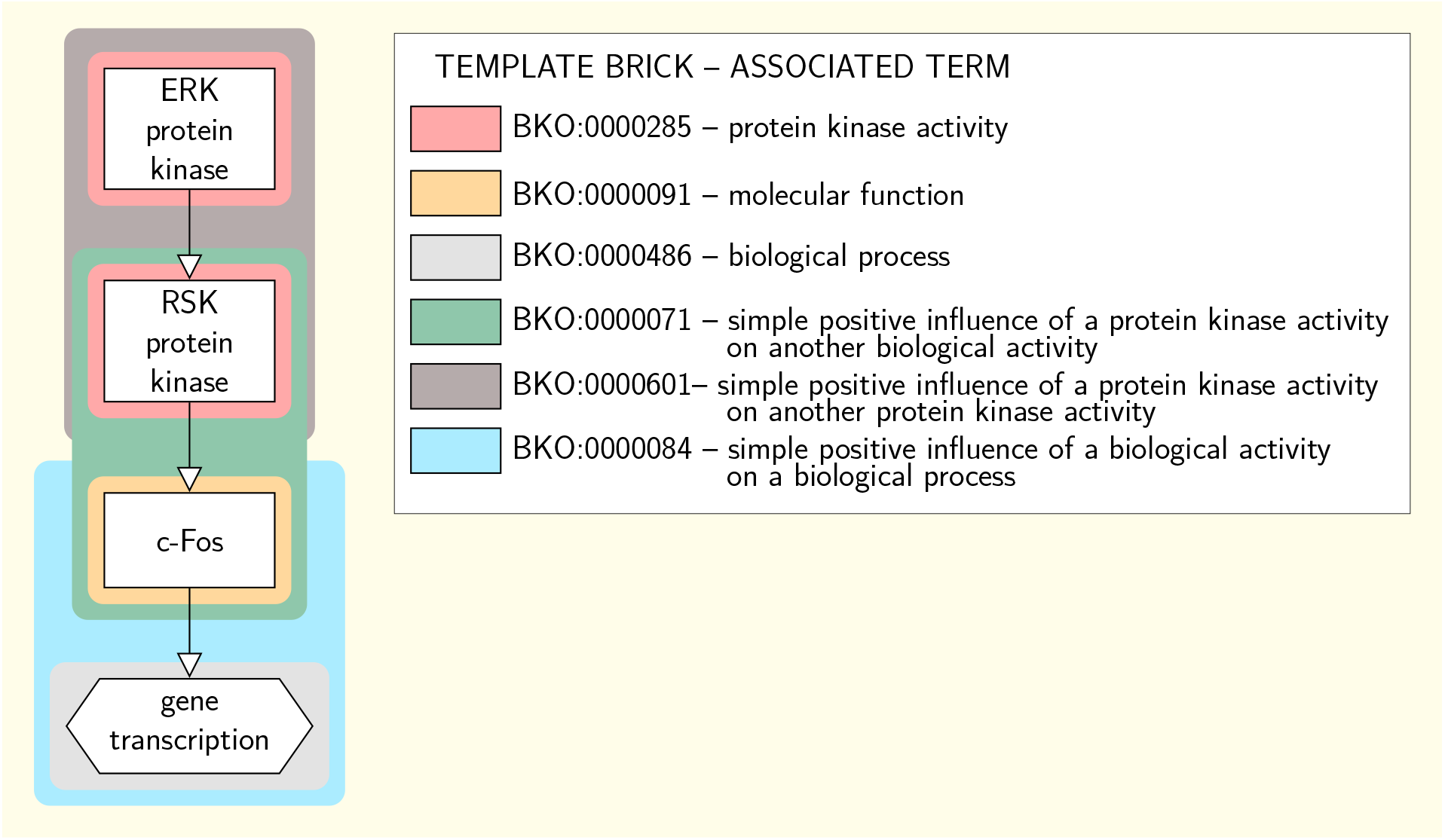
Excerpt of the SBGN AF map of the Insulin/IGF pathway-mitogen activated protein kinase/MAP kinase cascade annotated with terms using template bricks. This excerpt matches a part of the PD map of Figure 1. Colored boxes surround individual instance bricks matched by the template bricks given in the legend. The color of the surrounding box identifies the template brick the instance is matched by. Template bricks associated with the same terms as those associated with the PD template bricks of Figure 1 share the same color (e.g. the template brick BKO:0000285 associated with the term “protein kinase activity” is in pink, as is template brick BKO:0000287 in Figure 1 which is associated with the same term). This map is annotated with some terms that do not appear in the annotation of the PD map of Figure 1, indicating that those terms do not have any PD representation (e.g. the “simple positive influence of a protein kinase on another protein kinase”).

**Figure 6:**
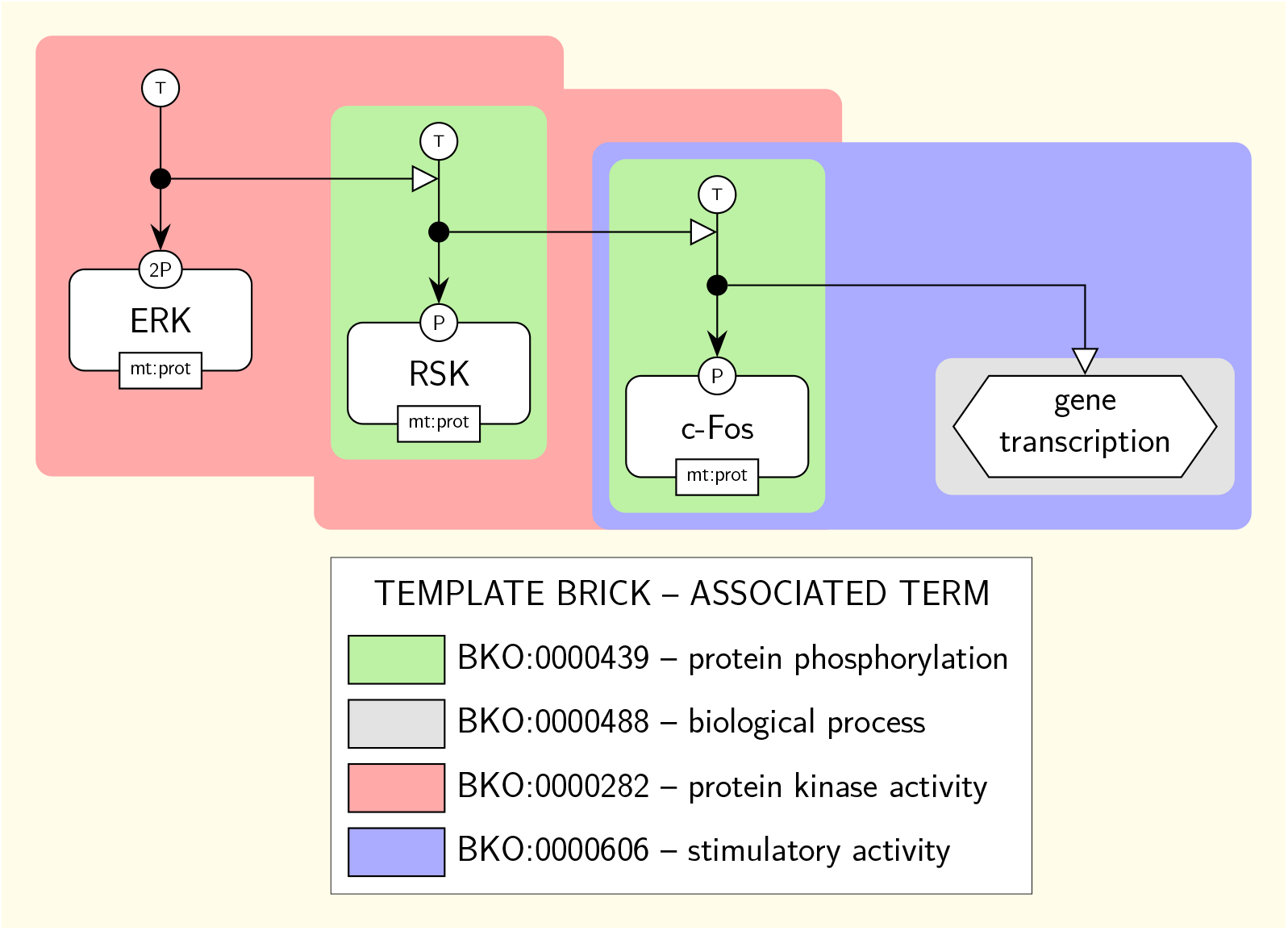
Excerpt of the SBGN ER map of the Insulin/IGF pathway-mitogen activated protein kinase/MAP kinase cascade annotated with terms using template bricks. This excerpt matches a part of the PD map of Figure 1. Colored boxes surround individual instance bricks matched by the template bricks given in the legend. The color of the surrounding box identifies the template brick the instance is matched by. Template bricks associated with the same terms as those associated with the PD template bricks of Figure 1 share the same color (e.g. the template brick BKO:0000285 associated with the term “protein kinase activity” is in pink, as is template brick BKO:0000287 in Figure 1 which is associated with the same term). Template brick BKO:0000287 in pink associated with the term “protein kinase activity” (GO:0004672) is an example of template brick associated with the term it represents using the *broad* relation.

The AF and ER template bricks are associated with the terms they represent in BKO using the relations described above for PD and two additional relations. The first of these relations, labeled synonym, is specific to AF template bricks. It is used when the template brick fully represents a *synonym* of the term that is not present in the ontology (some GO terms have synonyms that are not included in the ontology as terms, although they are explicitly associated with the main term). For example, the term “phosphoprotein phosphatase activity” (GO:0004721) has a template brick associated with it using the main relation (BKO:0000231) and two template bricks associated with it using the *synonym* relation: one for its *synonym* “protein phosphohydrolase” (BKO:0000236) and another for its *synonym* “protein phosphatase” (BKO:0000237). The second relation is labeled broad. It is used when the template brick represents a concept that is broader than the one described by the term and that is not described by any other term of the ontology. For example, the ER template brick representing the term “protein kinase activity” (GO:0004672), represented in pink in Figure 6 (BKO:0000282), actually represents a stimulation of a phosphorylation rather than a catalysis. Hence it represents a concept that is broader than the one described by the term (a catalysis being only a kind of stimulation) and is associated with the term using the *broad* relation.

### 2.4 Implementation and availability

BKO can be browsed and downloaded at sbgnbricks.org, and is registered at BioPortal (bioportal.bioontology.org/ontologies/BKO). As of June 2020, BKO contains 178 terms (42 imported from GO, 45 from SBO, and 91 newly defined), 476 template bricks (248 for PD, 70 for AF and 158 for ER), and 32 categories. All template bricks can also be freely downloaded in the form of PNG images or SBGN-ML files.

### 2.5 Evaluating the completeness of the ontology

To evaluate the completeness of our set of terms and template bricks with respect to the description of the main concepts found in biomolecular networks, we identified the set of instance bricks making up the PD maps of two databases: the Atlas of Cancer Signalling Network [17] (13 maps) and the PANTHER database [32, 22] (174 maps). To this end, for all maps, we identified all instances matching each PD template brick (see Methods for more details), resulting in zero or more instances for each template brick. Overall, we obtained 15708 distinct instances of BKO bricks for the ACSN, and 6339 for the PANTHER database. All processes and activities from both databases were matched by at least one template brick. This was to be expected since our set of template bricks covers all ways to represent the general kind of processes (irreversible/reversible processes) and activities (modulatory activity and binding activity) in PD. The number of instances matched by each term of the ontology for each of the two databases is given in Supplementary Table S2. To further evaluate the completeness of our ontology, we investigated those instance bricks that were matched by at least one of the 30 template bricks representing generic processes or activities (e.g. an irreversible process, a catalytic activity), without being matched by template bricks representing more specific processes or activities, and that are subclasses of the former in the ontology (e.g. a protein phosphorylation, a protein kinase activity). Those instances could reveal that a particular term or template brick is lacking in the ontology. The list of the 30 template bricks representing generic concepts is given in Supplementary Table S3. Across the two databases, there were 6396 of such instances, representing 28% of the total number of instances. For each instance, we identified the nature of the process or the activity it represented manually (see Supplementary Table S4-5). Based on this analysis, we subsequently categorized all instances in one of six categories. Results are given in Table 1, and examples in Supplementary Figure S3. Approximately 56% of these instances could not be matched to more specific template bricks for reasons independent to the ontology: we found that 26% were misrepresentations (e.g. typos, incomplete processes or invalid SBGN representations such as processes consuming or producing phenotypes), 30% represented processes whose nature was implicitly given by the labels of its participants (e.g. a protein truncation leading to the production of proteins with new labels) and less than 1% represented processes whose nature we could not identify. Another 37% of the instances represented a modulatory activity involving a process whose nature corresponded to a term of the ontology but that itself did not correspond to any specific term (e.g. the stimulation of a dissociation). Terms describing such modulations we not integrated to the ontology as they are not sufficiently relevant with regard to bio-molecular activities. Another 1% of the instances involved two processes of different nature represented using only one SBGN process (e.g. a mix between a dissociation and a phosphorylation). For each such instance, both processes taken individually corresponded to a term of the ontology. However, template bricks representing such hybrid processes were not integrated to the ontology as these processes are more conventionally represented by a sequence of two more elementary processes. Finally, 6% of the instances involved a process whose nature did not correspond to any specific term of the ontology. Terms describing these processes were not integrated to the ontology either because these processes are not clearly identified in regard to biology (non standard values in state variables such as “?” or “*” (2.5%); production from an empty set (0.08%); methylation of a nucleic acid feature of unspecified nature (0.06%); creation of asRNAs (1.6%)) or because their bio-molecular mechanisms involves a sequence of more elementary processes (GTP/GDP exchange process (1.8%)). Overall, our analysis showed that BKO is complete enough to accurately represent all types of processes and activities found in the maps of the two databases. Additionally, it revealed that BKO contains terms and template bricks that do not match any instances in both databases, suggesting that some terms and bricks may be less relevant than others from a practical point of view. However they participate to the completeness of the ontology from a theoretical point of view and may be valuable for future methodological developments and analyses, which may be assessed in the future by repeating the present analysis on maps found in additional databases.

**Table 1:**
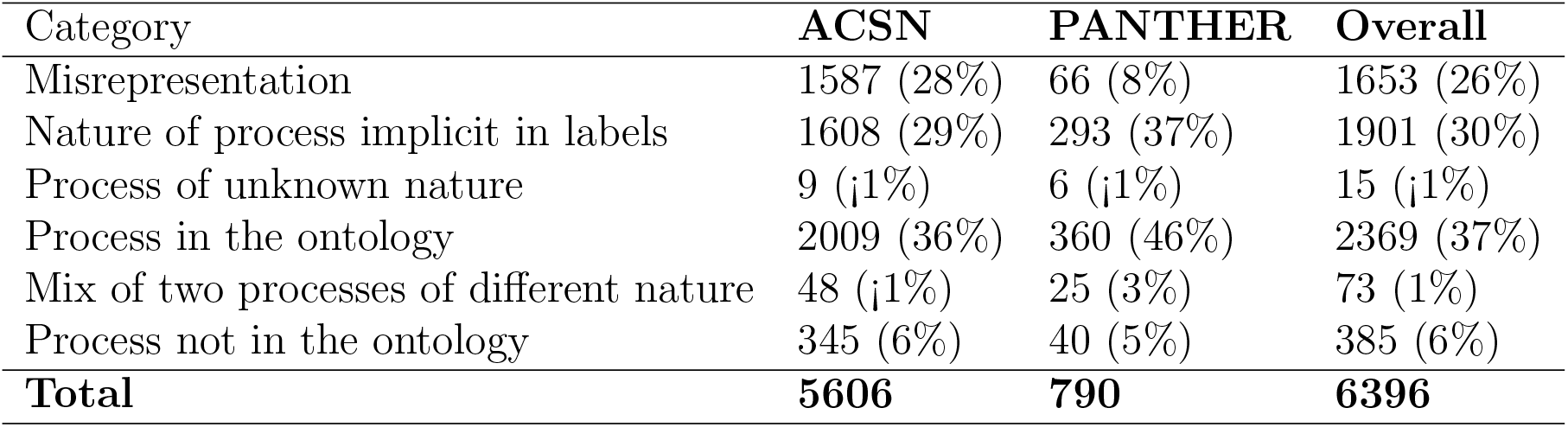
Classification of all instances of the ACSN and PANTHER databases that were only matched by template bricks representing generic concepts. All bricks that were matched by one of 30 template bricks representing generic concepts (e.g. an irreversible process, a catalytic activity) without being matched by template bricks representing more specific concepts (e.g. a protein phosphorylation, a protein kinase activity) were classified in one of the following six categories. **Misrepresentation**: the process or activity is badly represented (e.g. it involves a phenotype as a reactant or product which is not valid SBGN). **Nature of process implicit in labels**: the nature of the process can solely be identified by analyzing the labels of the participants of the process (e.g. the truncation of a protein, where a protein is consumed to give two or more proteins with new names). **Process of unknown nature**: we could not identify the nature of the represented process or the activity. **Process in the ontology**: this category implies only modulatory activities whose modulated process is an instance of a template brick that belongs to the ontology (e.g. stimulation of dissociation). **Mix of two processes of unknown nature**: two processes are represented using only one SBGN process (e.g. a dissociation and a dephosphorylation). **Process not in the ontology**: the process represented in the brick is not an instance of any template brick of the ontology.

## 3 Discussion

### 3.1 Generalization to other languages

While we specifically built template bricks expressed using the three SBGN languages, our approach was designed to be universally applicable to any language used to describe molecular networks, for example, BioPAX [9] or SBML [14, 13]. On a theoretical point of view, the definitions of a template brick and an instance brick, and their integration in the ontology under the form of a class or an individual, can be generalized to any such language. In addition, we implemented BKO so that it is language-agnostic and that template bricks expressed in any language can easily be added to it. Building sets of template bricks expressed in various standard formats and integrating them in BKO alongside the SBGN bricks would allow for a better alignment of these standards, which could constitute a solid conceptual base for the development of new conversion tools.

### 3.2 Updating BKO

We intend to update BKO following the respective updates of the SBO and the GO ontologies, and of the SBGN standard. We expect that these updates will consist mostly of imports into BKO of new terms of SBO and GO and the addition of new template bricks associated with those terms, as well as template bricks represented using new versions of the SBGN languages.

### 3.3 Expected methodological impact

The first set of SBGN bricks introduced in Junker *et. al* [15] has been used as a teaching aid and as an SBGN learning resource (Learning SBGN, sbgn.org/learning. It also served as a basis for the development of a number of methods and tools related to the SBGN standard. Its support advanced the development of SBGN-specific automatic layout algorithms in the yEd Graph Editor via the SBGN Palette and SBGN Layout functionalities [29]. Also, it became clear that there was a promise for conversion between the conceptually different SBGN languages. A first alignment of template bricks was shaped in the SBGN Bricks Dictionary (available at sbgnbricks.sourceforge.net/sbgnbricks_dictionary.html) and was used for the transformation of PD maps to AF maps [36]. It was later used for the development of the PD2AF conversion rules and the PD2AF tool (www.pd2af.org) designed for the same type of transformation. The notion of bricks also contributed to the development of the conversion rules of the STON tool that allows storing and querying PD maps in a Neo4j Graph Database [33], of the conversion rules of the cd2sbgnml bidirectional converter [2] between the CellDesigner format [11] and PD maps expressed in the SBGN-ML format [3], and of the conversion rules of ySBGN (github.com/sbgn/ySBGN), a bidirectional converter between yEd (www.yworks.com/products/yed) GraphML format and the SBGN-ML format. Recently this approach was also used as a basis for the development of the ModelBricks project [7] (modelbricks.org), which enables the on demand generation of executable models from PD maps. We expect that the large extension of the set of template bricks and its formal representation in BKO will lead to the development of new methods and tools. In particular, the association of template bricks with terms precisely describing the concepts encountered in molecular networks paves the way to the development of tools based on the annotation of SBGN maps. Such an annotation is for example a prerequisite for the semantic validation of maps, or for their merging. In this regard, BKO could be used to improve map curation workflows and tools, by for example allowing automatic detection of misrepresentations or repetitions. In addition, extending BKO with sets of bricks expressed in other standard languages such as BioPAX will allow the development of new converters grounded in a formal framework. Such an extension will also open up the possibility to integrate the annotation process we used to analyze the ACSN and PANTHER databases into more systematic workflows [19] while using such model repositories as BioModels [4], Physiome Model Repository [28] or Reactome [10]. Formally represented building blocks of SBGN maps will also enable integration of graphical information into sophisticated model retrieval tools such as MaSyMoS [12].

## 4 Methods

### 4.1 Representation of template bricks and matching rules

The template bricks are generic patterns that may be used to generate or match specific instances. They are represented using the glyphs of the three SBGN languages and additional textual constructs that allow expressing generality or repetition, for example. The definition of these elements stemmed from a balance between three requirements: (i) they should allow describing generic patterns that could be used for pattern matching as well as for template-based construction; (ii) they should not distort the meaning of the SBGN glyphs specified in the SBGN specifications; and (iii), they should remain as simple as possible so they could be easily understood and used. We describe these elements and the rules defining how they are used to match real instances hereafter.

#### 4.1.1 Matching of glyphs and completion rules

##### Matching of glyphs

The template bricks are represented using the glyphs defined by the SBGN standard. When used in a template brick, we call such a glyph a template glyph. For a template brick to match a given instance, all the template glyphs it is formed of must match a different glyph of the instance. The matching rules for glyphs are as follows. In general, a template glyph matches any instance of its SBGN counterpart, and is represented the same way (i.e. using the same shape). The only template glyph for which this matching rule does not apply is the macromolecule template glyph (in PD). Indeed, the macromolecule glyph is used to represent a pool of macromolecules (e.g. proteins, polysaccharides), and the specific nature of the macromolecules may be indicated by decorating the glyph with a unit of information defining a material type (e.g. “mt:prot” for proteins, “mt:psac” for polysaccharides). However, in practice, the macromolecule glyph is mostly used without such a decoration to represent a pool of proteins. Therefore, we defined the following specific matching rule for the macromolecule template glyph: it matches an instance of its SBGN counterpart only if this instance is not decorated by a unit of information defining a material type other than “mt:prot”.

We defined additional matching rules for some template glyphs, which allows defining more generic patterns. A reduced number of template glyphs match instances of additional SBGN glyphs that are not their counterpart, in a manner consistent with their ontological meaning. For example, the modulation template glyph matches any instance of its SBGN counterpart but also any instance of all other SBGN glyphs representing other types of modulations, since this modulation is generic and can be put at the top of the ontology describing the different types of modulation. Figure 7 shows this ontology (for PD). Each template glyph of the ontology matches any instance of its SBGN counterpart, as well as any instance of the SBGN counterparts of all its descendants. Hence the stimulation template glyph will match any instance of the SBGN stimulation glyph, but also of the SBGN necessary stimulation and catalysis glyphs. Analogously, the modulation template glyph will match any instance of the SBGN modulation, inhibition, stimulation, necessary stimulation, and catalysis glyphs. We consider the same type of matching for all template glyphs representing generic concepts, i.e. glyphs representing processes, entity pools and subunits of complexes in PD, and glyphs representing modulations or influences in ER and AF, respectively. In particular, the unspecified entity template glyph (PD) matches any instance of an SBGN glyph representing an entity pool, such as a macromolecule or a simple chemical glyph, and the process template glyph (also PD) matches any instance of an SBGN glyph representing a stoichiometric process (that is all glyphs representing processes but the phenotype glyph). The layout of the glyphs is generally not considered in the matching, except in the following cases describing qualitative spatial relationships: entity pool nodes and activities inside compartments, unit of information and state variables decorating glyphs, and subunits inside complexes.

**Figure 7:**
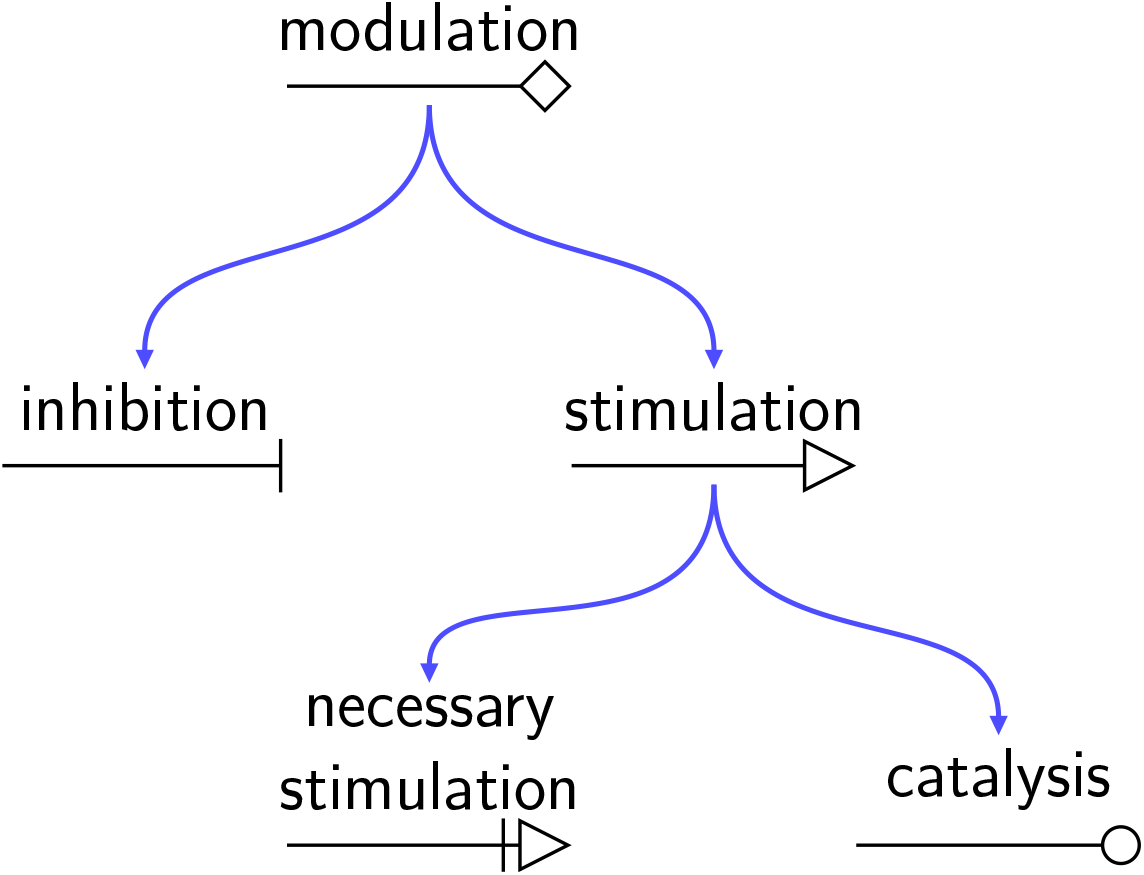
Ontology organizing the template glyphs representing modulations (PD). Each template glyph matches any instance of its SBGN counterpart as well as any instance of the SBGN counterparts of all its descendants.

##### Completion rules

The template bricks only represent the core of each concept. Instances in real maps may include more glyphs than those represented in the templates. For example, a PD representation of the phosphorylation of a given protein may include the consumption of ATP and production of ADP, which are not represented in the corresponding template brick (BKO:0000440). These glyphs will therefore not be matched by the template brick. However, they represent parts of the phosphorylation process and should be included in the matching result. For this reason, we defined rules for the completion of matched instances. They include rules for the completion of glyphs with auxiliary units, with compartments, and with process participants. These rules are applied recursively.

- *Auxiliary units*: Any instance glyph matched by the template brick is completed with all the instance auxiliary units decorating it that are not subunits (inside complexes). This includes units of information and state variables for entity pool nodes, for example.
- *Compartments*: Any instance glyph matched by the template brick is completed with the instance compartment containing it, if any. Containment in compartments is only defined for entity pool nodes and activities.
- *Process participants*: Any instance of a process glyph matched by the template is completed with all instance glyphs that are linked to this process by a consumption or a productuction arc.

#### 4.1.2 Additional textual constructs

We defined additional constructs that allow expressing generality or repetition, for example. These constructs are always contained in the labels of the template glyphs, and are expressed using specific symbols that serve as delimiters (“$”, “[“, “]”, “*{*“, “*}*” and “!”). The first constructs we introduce are strictly related to the matching of labels, while the others are related to the matching of topologies that cannot be expressed using only the set of template glyphs.

##### Labels

- *Literal* : A string not enclosed by “$” delimiters matches itself. For example, in Figure 8 panel A, literal “ERK” matches itself (top) but not “ERK1” (center) nor “c-Fos” (bottom).
- *Variable string* : A string enclosed by the “$” delimiters, such as “$NAME$”, matches any non-empty string. The enclosed string, here “NAME”, acts as a variable name. Hence, all occurrences of “$NAME$” in a template brick must match the same string. For example in Figure 8 panel B, “$UNSPEC$” matches “ERK” (top) and “$COMP$” matches “nucleus” (top and bottom). However, “$UNSPEC$” may not match both “ERK-cyt” and “ERK-nuc” at the same time (bottom).
- *Optional string* : A string enclosed by the “[“and “]” delimiters, such as “[CONTENT]”, is optional. Hence, “[CONTENT]” may match any string that may be matched by CONTENT, or the empty string. In Figure 8 panel C, “[@$RES$]” matches either the empty string (top), or “@S221” (bottom).
- *Disjunction*: The “—” character is used as a disjunction operator between two literals. Hence, a construct of the form “LITERAL 1—LITERAL 2” matches either “LITERAL 1” or “LITERAL 2”. For example, in Figure 8 panel D, “cell—CELL” matches either “cell” (top) or “CELL” (bottom).

**Figure 8:**
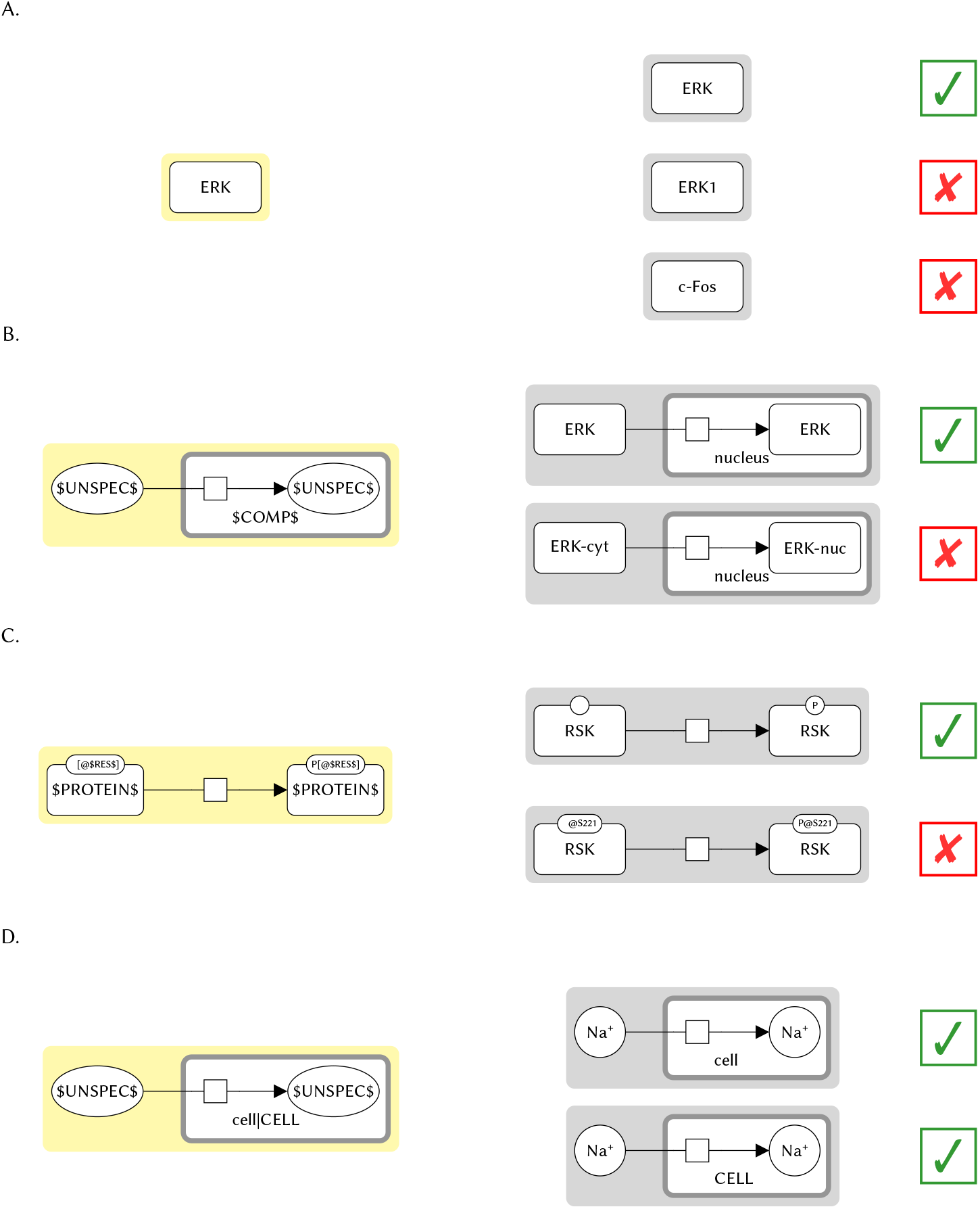
Textual constructs for label matching. Template bricks are represented on the left with a yellow background (one per construct), and instances are represented on the right of each template brick with a gray background. The instances matched by the template brick are indicated with a check mark, and those not matched with a cross. **A**. Literal. “ERK” matches itself (top) but not “ERK1” (center) nor “c-Fos” (bottom). **B**. Variable string. The template brick (BKO:0000484) represents the term “translocation reaction” (SBO:0000185). “$UNSPEC$” matches “ERK” (top) and “$COMP$” matches “nucleus” (top and bottom). “$UNSPEC$” may not match both “ERK-cyt” and “ERK-nuc” at the same time (bottom). **C**. Optional string. The template brick (BKO:0000440) represents the term “protein phosphorylation” (BKO:0000438). “[@$RES$]” matches either the empty string (top) or “@S221” (bottom). Likewise, “P[@$RES$]” matches either “P” (top), or “P@S221” (bottom). **D**. Disjunction. The template brick (BKO:0000489) represents the term “transcellular membrane influx reaction” (SBO:0000587). “cell—CELL” matches either “cell” (top) or “CELL” (bottom).

##### Topology

- *Repetition*: The ““and”” delimiters are used to express repetition. They are always put at both ends of the label of a glyph, and apply to the whole glyph. A glyph with a label of the form “CONTENT” may match zero or more instance glyphs that may be matched by the template glyph with the ““and”” delimiters removed. The match is greedy. In the current set of template bricks, this construct is used to explicitly represent additional reactants or products of processes, additional subunits of complexes (PD), or additional interaction participants (ER). For example in Figure 9 panel A, the template glyph with label “SUBSTRATE 1” matches one of the three reactants of the process of the instance (right), while the template glyph with label “$SUBSTRATE_N$” (left) matches the other two (right). Additionally, the repetition construct acts as a negative lookup, which allows expressing exclusivity. The match will not be successful if the instance contains a glyph not matched by any template glyph of the template and that has the same role as the template glyph with the repetition construct (e.g. is also a reactant of a process, or a subunit of a complex). For example, in Figure 9 panel A, the template brick (left) will not match any instance representing a process consuming or producing an entity pool that is not a simple chemical.
- *Absence*: The “!” delimiters are used to match the absence of a glyph (negative lookup). They are always put at both ends of the label of a glyph, and apply to the whole glyph. A glyph with a label of the form “!CONTENT!” matches the absence of any glyph that may be matched by the template glyph with the “!” delimiters removed (i.e., such an instance must not be present for the match to be successful). In the current set of template bricks, this construct is used only for the template brick representing a “passive transport” (SBO:0000658) which is defined as a transport with no consumption of ATP. This template brick is represented in Figure 9, panel B (left). The process of the template consumes a simple chemical with label “!ATP!” and produces one with label “!ADP!”, indicating that it will not match any instance whose process consumes a simple chemical with label “ATP” or produces one with label “ADP”. It is the case for the instance, hence it is not matched (right).

**Figure 9:**
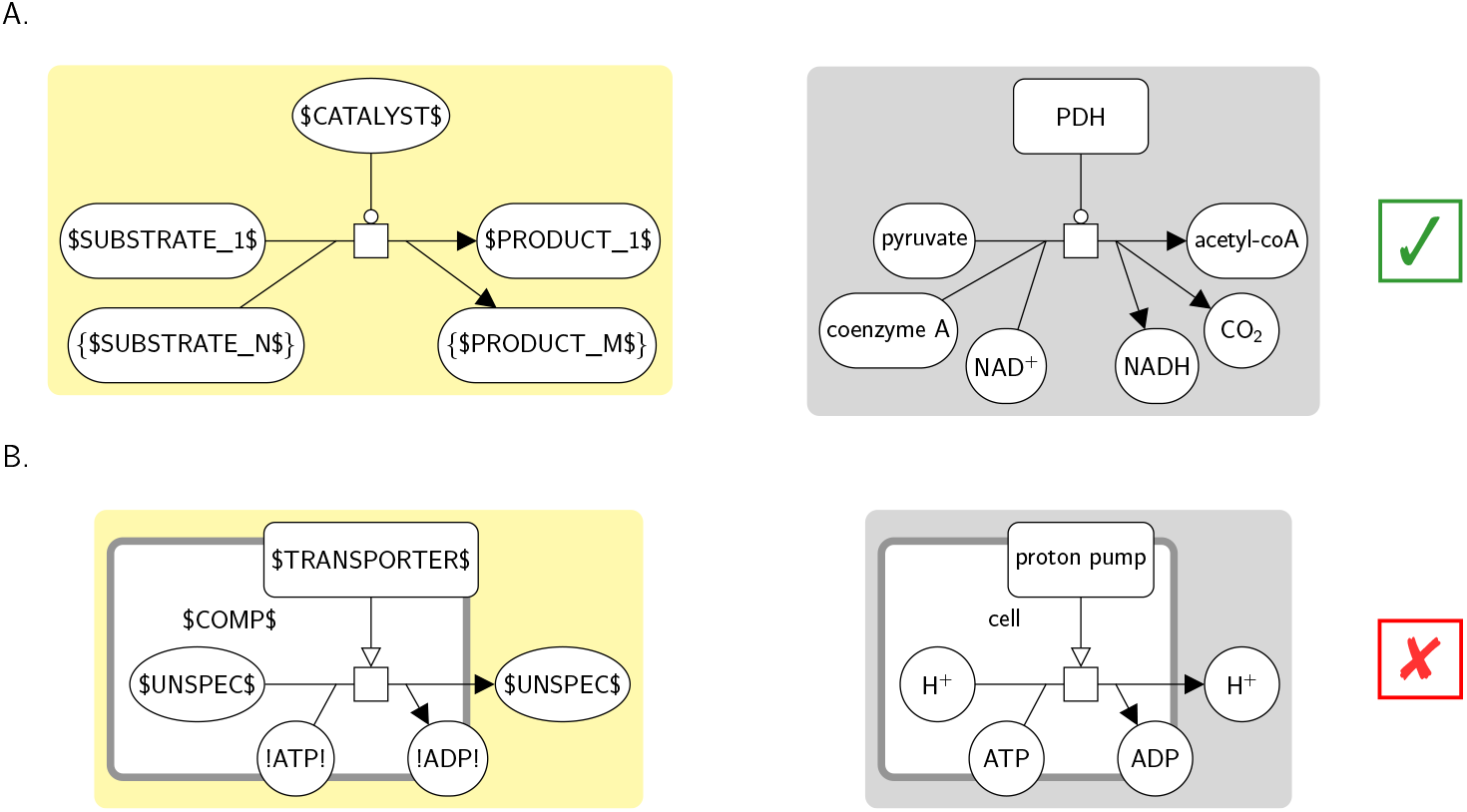
Textual constructs for the matching of topologies that cannot be expressed with the template glyphs alone. Template bricks are represented on the left with a yellow background (one per construct), and instances are represented on the right of each template brick with a gray background. The instances matched by the template brick are indicated with a check mark, and those not matched with a cross. **A**. Repetition. The template brick (BKO:0000197) represents the term “metabolic catalytic activity” (BKO:0000196). The template glyph with label $SUBSTRATE_N$ may match zero or more simple chemical glyphs consumed by the process, here two of the three reactants. Likewise the template glyph with label $PRODUCT M$ may match zero or more simple chemicals produced by the process, here two of the three products. **B**. Absence. The template brick (BKO:0000492) represents the term “passive transport” (SBO:0000658). The template simple chemical with label “!ATP!” will match the absence of a simple chemical reactant with label “ATP”, and the one with label “!ADP!” the absence of a simple chemical product with label “ADP”. Hence the instance is not matched, because it consumes and produces such simple chemicals.

### 4.2 Construction of the set of template bricks and BKO

The new set of 476 template bricks and BKO were built in a three steps process. First, we enumarated all concepts described in SBO and GO that could be represented using SBGN in a generic way. We built at least one template brick for each of these terms, and integrated the newly built template bricks and their corresponding terms in a new ontology (BKO). Then we completed BKO with terms and template bricks describing and representing concepts that were not present in SBO or GO but that could be represented using SBGN in a generic way. These concepts were either more specific than those of SBO and GO (e.g. a protein phosphorylation specifies a phosphorylation (SBO)) or more generic (e.g. a stimulatory activity generalizes a catalytic activity (GO)). Finally we completed BKO by analyzing real SBGN maps taken from the ACSN and PANTHER databases.

BKO was built using Protege [24] and the owlready2 Python library [20]. Terms and template bricks were added in the ontology as classes, and the different categories as individuals of a unique category class. Associations between template bricks and terms/categories were added in the ontology as class annotations. Among the 177 terms of the ontology, 42 terms were imported from GO [1, 5], and another 45 terms were imported from SBO [6].

Finally, the template bricks were represented in SBGN using the sbgntikz library [25] and SBGN-ED [8].

### 4.3 Pattern matching in the maps of the ACSN and PANTHER databases

Each map of the ACSN [17] and PANTHER [32, 22] databases was converted from the CellDe-signer format (celldesigner.org/features.html) to the SBGN-ML format [3] using the cd2sbgnml converter [2], with a posteriori addition of active and inactive state variables to macromolecules and complexes, and of units of information with label “ct:mRNA” (resp. “ct:gene”) to mRNAs (resp. genes). Each map was then stored in a Neo4j database (neo4j.com) using stonpy v0.1.x (github.com/adrienrougny/stonpy), a new Python 3.x version of STON [33] specifically developed for easing the identification of instance bricks in SBGN maps. Each PD template brick was then expressed as a Cypher query (the query language for Neo4j graph databases) following the matching rules introduced earlier, and was run against the database containing the maps. Results of each query were then transformed into individual SBGN maps (instances) and stored in the SBGN-ML format using stonpy.

## 5 Conclusion

The concept of SBGN bricks has been employed for the development of multiple tools including format converters and template-based functionalities in editors. This shaped a demand in an extended and better-organised set of SBGN bricks for providing better support for such tools. We introduce the BricKs Ontology (BKO) that includes a new set of 476 bricks hierarchically organised. BKO is downloadable and browsable online at sbgnbricks.org, and is available under a CC BY 4.0 license. The completeness of this updated set of bricks was evaluated by checking them against the maps of the ACSN and PANTHER pathway databases. All processes and activities could be accurately identified and annotated, suggesting that BKO may be a valuable resource for the description of recurring concepts in molecular networks. We expect that the new extended set of bricks and the new semantic layer brought by BKO will enable the development of new methods and tools for solving long-standing problems that require semantic analysis of SBGN maps, such as quality check, network comparison, merging of maps or the transformation between SBGN Process Description and Activity Flow languages. Moreover, we intend to extend the BKO with bricks built using other standard formats, which will allow a better alignment of these formats and the development of converters grounded in a formal conceptual framework, thus contributing to the collaborative evolvement of systems biology approaches and avoiding redundant efforts.

## Acknowledgements

This work was conducted using the Prot égé resource, which is supported by grant GM10331601 from the National Institute of General Medical Sciences of the United States National Institutes of Health.

