## Supplementary Material for "SBGN Bricks Ontology as a tool to describe recurring concepts in molecular networks"

October 28, 2020

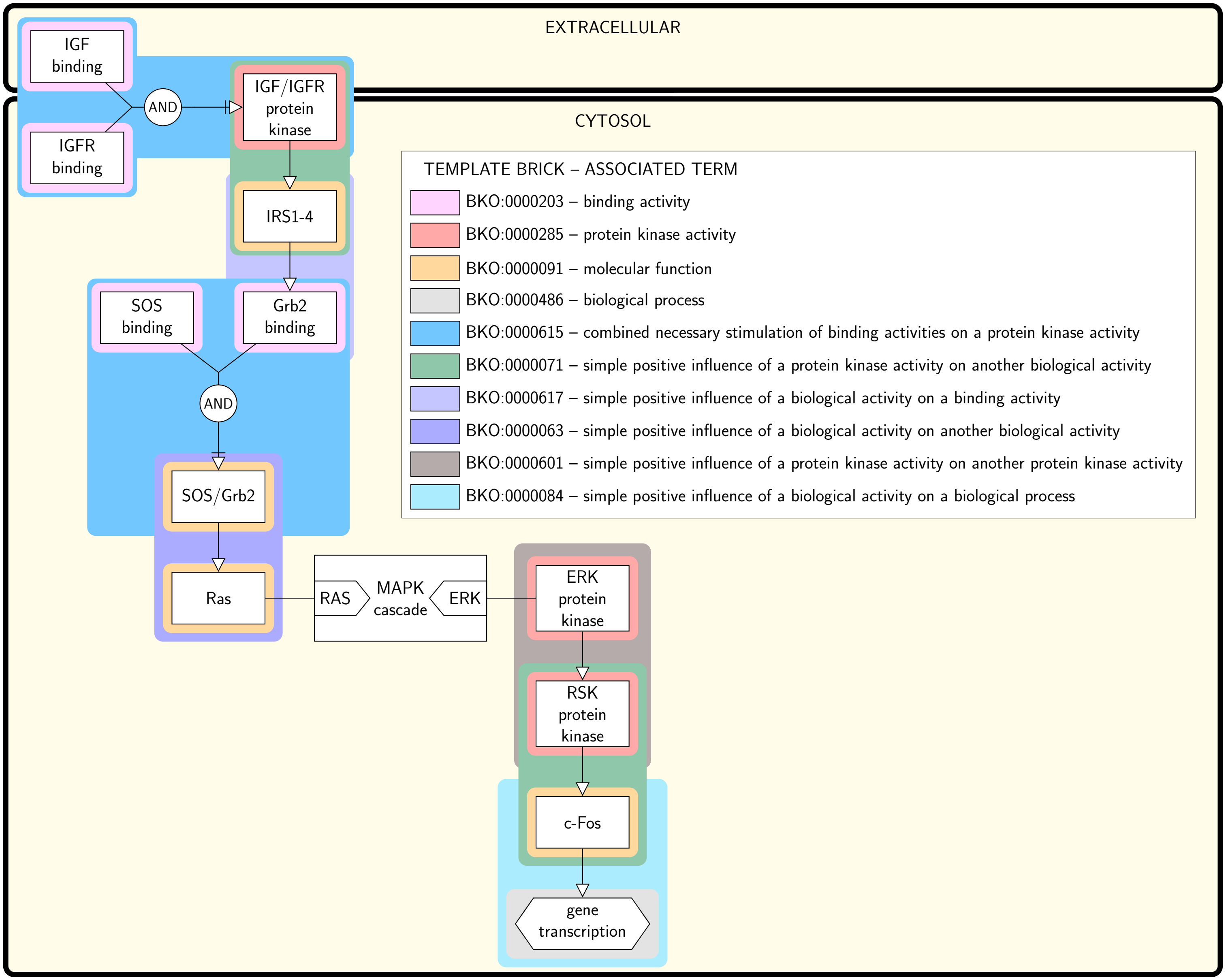


Figure S1: **SBGN AF map of the Insulin/IGF pathway-mitogen activated protein kinase/MAP kinase cascade annotated with terms using template bricks.** This map matches the PD map of Figure 1. Colored boxes surround individual instance bricks matched by the template bricks given in the legend. The color of the surrounding box identifies the template brick the instance is matched by. Template bricks associated with the same terms as those associated with the PD template bricks of Figure 1 share the same color (e.g. the template brick BKO:0000285 associated with the term "protein kinase activity" is in pink, as is template brick BKO:0000287 in Figure 1 which is associated with the same term). This map is annotated with some terms that do not appear in the annotation of the PD map of Figure 1, indicating that those terms do not have any PD representation (e.g. the "simple positive influence of a protein kinase on another protein kinase").

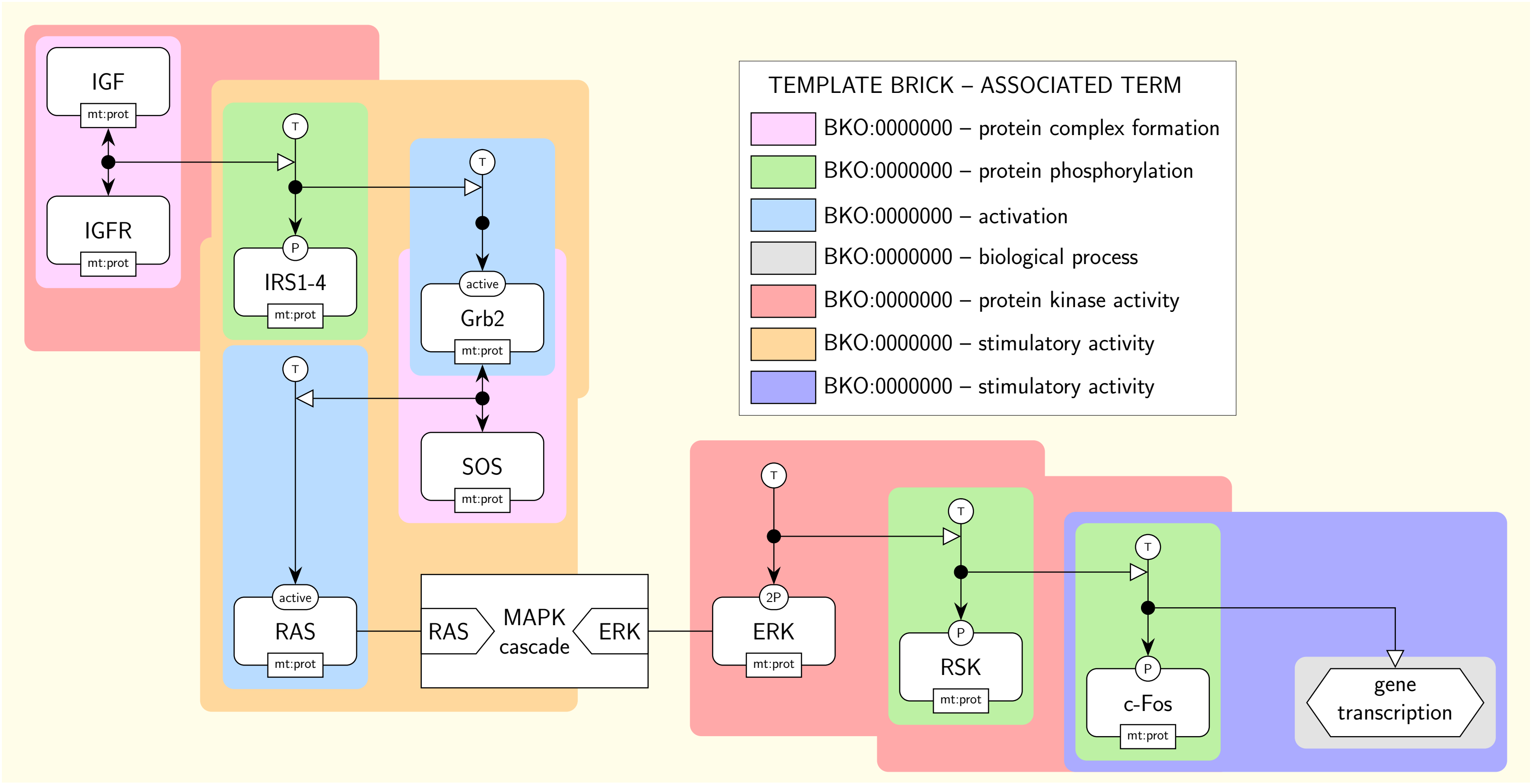


Figure S2: **SBGN ER map of the Insulin/IGF pathway-mitogen activated protein kinase/MAP kinase cascade annotated with terms using template bricks.** This map matches the PD map of Figure 1. Colored boxes surround individual instance bricks matched by the template bricks given in the legend. The color of the surrounding box identifies the template brick the instance is matched by. Template bricks associated with the same terms as those associated with the PD template bricks of Figure 1 share the same color (e.g. the template brick BKO:0000285 associated with the term “protein kinase activity” is in pink, as is template brick BKO:0000287 in Figure 1 which is associated with the same term).

Table S1: **Alignment of PD, AF and ER template bricks associated with the terms describing common concepts encountered in molecular networks.**

| Term | PD | AF | ER |
|---|---|---|---|
| SBO:0000177 non-covalent binding - GO:0005488 binding | 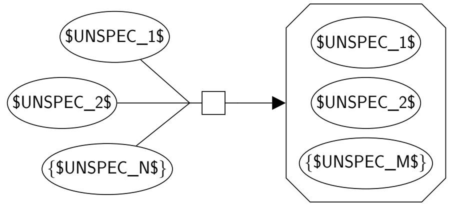  BKO:0000204 non-covalent binding (PD narrow 1) | 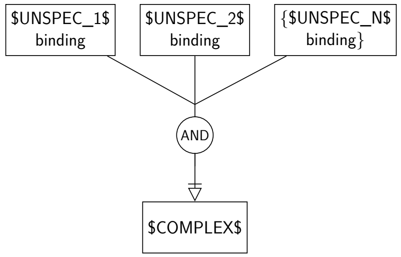  BKO:0000203 binding (AF main) - BKO:0000042 combined necessary stimulation of binding activities on another biological activity (AF main) | 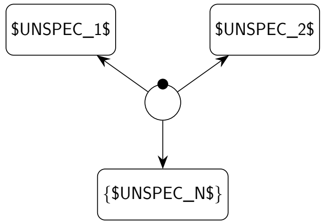  BKO:0000207 non-covalent binding (ER main) |
| BKO:0000221 protein multimerisation | 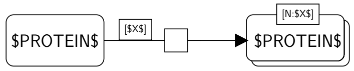  BKO:0000223 protein multimerisation (PD main) | 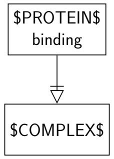  BKO:0000203 binding (AF main) - BKO:0000059 simple necessary stimulation of a binding activity on another biological activity (AF main) | 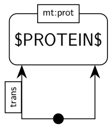  BKO:0000222 protein multimerisation (ER main) |
| SBO:0000180 dissociation | 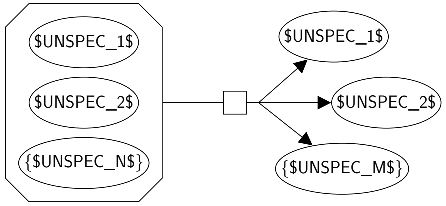  | Not applicable | Not applicable |

| | | | |
|---|---|---|---|
| | BKO:0000170 dissociation (PD narrow 1) | | |
| BKO:0000438 protein phosphorylation | 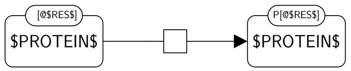<br>BKO:0000440 protein phosphorylation (PD narrow 1) | Not applicable | 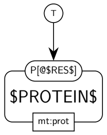<br>BKO:0000439 protein phosphorylation (ER main) |
| GO:0004672 protein kinase activity | 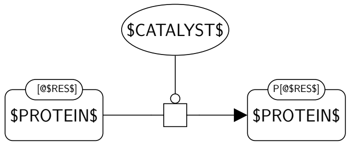<br>BKO:0000287 protein kinase activity (PD narrow 1) | 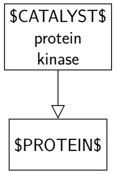<br>BKO:0000285 protein kinase activity (AF main) - BKO:0000071 simple positive influence of a protein kinase activity on another biological activity (AF main) | 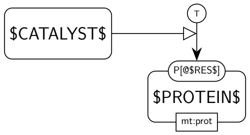<br>BKO:0000282 protein kinase activity (ER broad 1) |
| BKO:0000375 protein dephosphorylation | 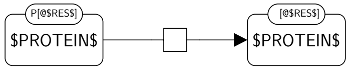<br>BKO:0000370 protein dephosphorylation (PD narrow 1) | Not applicable | 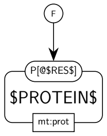<br>BKO:0000369 protein dephosphorylation (ER main) |

| | | | |
|---|---|---|---|
| GO:0004721 phosphoprotein phosphatase activity | 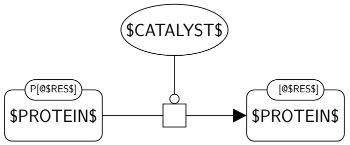  BKO:0000233 phosphoprotein phosphatase activity (PD narrow 1) | 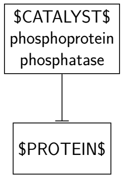  BKO:0000068 simple negative influence of a phosphoprotein phosphatase activity on another biological activity (AF main) | 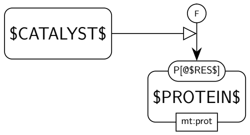  BKO:0000228 phosphoprotein phosphatase activity (ER broad 1) |
| SBO:0000656 activation | 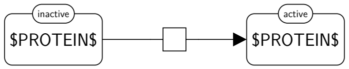  BKO:0000498 activation (PD narrow 1) | Not applicable | 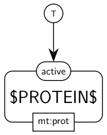  BKO:0000495 activation (ER main) |
| SBO:0000208 deactivation | 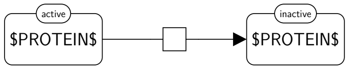  BKO:0000211 deactivation (PD narrow 1) | Not applicable | 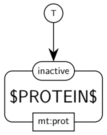  BKO:0000210 deactivation (ER main) |
| BKO:0000511 irreversible metabolic reaction | 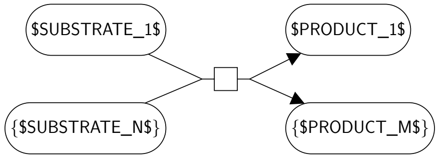  BKO:0000513 irreversible metabolic reaction (PD main) | Not applicable | Not applicable |

| | | | |
|---|---|---|---|
| BKO:0000514 reversible metabolic reaction | 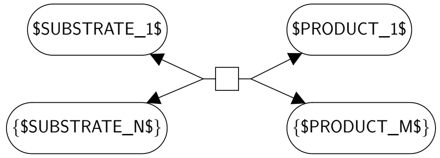<br>BKO:0000515 reversible metabolic reaction (PD main) | Not applicable | Not applicable |
| BKO:0000196 metabolic catalytic activity | 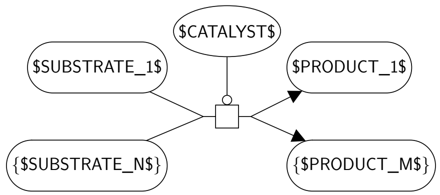<br>BKO:0000197 metabolic catalytic activity (PD narrow 1) | Not applicable | Not applicable |
| BKO:0000196 metabolic catalytic activity | 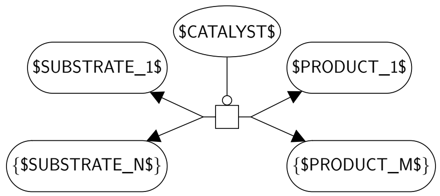<br>BKO:0000198 metabolic catalytic activity (PD narrow 2) | Not applicable | Not applicable |
| SBO:0000183 transcription | 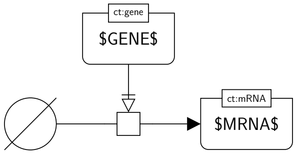 | 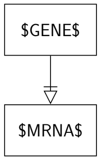 | 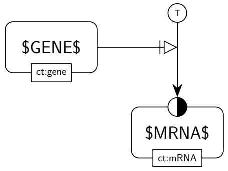 |

| | BKO:0000477 transcription (PD main) | BKO:0000061 simple necessary stimulation of a biological activity on another biological activity (AF main) | BKO:0000478 transcription (ER main) |
|---|---|---|---|
| GO:0001216 DNA-binding transcription activator activity | 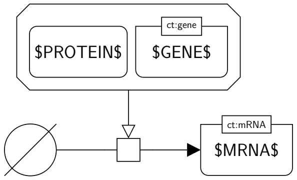<br>BKO:0000518 DNA-binding transcription activator activity (PD main) | 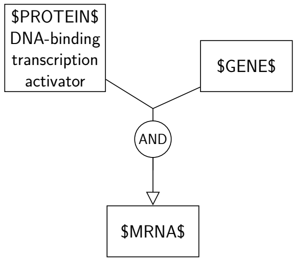<br>BKO:0000520 DNA-binding transcription activator activity (AF main) - BKO:0000055 combined positive influence of a DNA-binding transcription activator activity and a biological activity on another biological activity (AF main) | 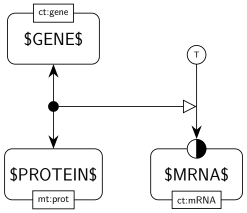<br>BKO:0000519 DNA-binding transcription activator activity (ER main) |
| SBO:0000184 translation | 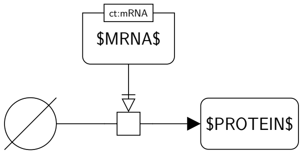<br>BKO:0000481 translation (PD main) | 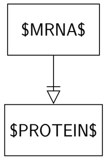<br>BKO:0000061 simple necessary stimulation of a biological activity on another biological activity (AF main) | 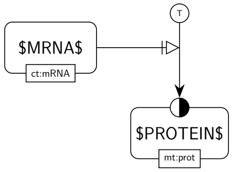<br>BKO:0000480 translation (ER main) |
| BKO:0000508 channel opening | 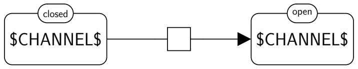<br>BKO:0000509 channel opening (PD main) | Not applicable | 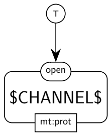<br>BKO:0000510 channel opening (ER main) |

| | | | |
|---|---|---|---|
| BKO:0000505 channel closing | 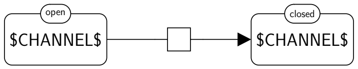 BKO:0000506 channel closing (PD main) | Not applicable BKO:0000507 channel closing (ER main) | 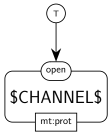 |
| SBO:0000185 translocation reaction | 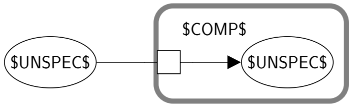 BKO:0000484 translocation reaction (PD narrow 2) | Not applicable | 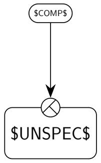 BKO:0000482 translocation reaction (ER main) |
| SBO:0000655 transport reaction - GO:0005215 transporter activity | 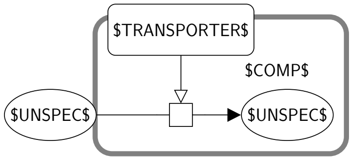 BKO:0000090 transport reaction (PD narrow 2) | 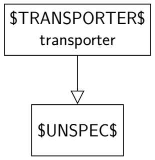 BKO:0000088 transporter activity (AF main) - BKO:0000073 simple positive influence of a transporter activity on another biological activity (AF main) | 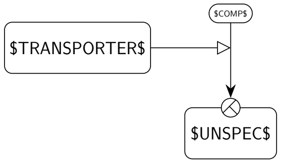 BKO:0000087 transport reaction (ER main) |

Table S2: **Number of instances matched by each term of the ontology in the ACSN and PANTHER databases.** The number in parantheses is the number of proper instances matched by the term, i.e. the number of instances that are matched by the term and by none of its descendants. Terms matching no instances in both databases are not shown.

| Term | ACSN | PANTHER | Overall |
|---|---|---|---|
| BKO:0000041 term | 15708 (0) | 6339 (0) | 22047 (0) |
| BKO:0000078 molecular function | 7865 (0) | 3165 (0) | 11030 (0) |
| BKO:0000008 modulatory activity | 6275 (66) | 2638 (0) | 8913 (66) |
| BKO:0000005 stimulatory activity | 5207 (515) | 2470 (54) | 7677 (569) |
| BKO:0000004 catalytic activity | 4692 (2547) | 2416 (517) | 7108 (3064) |
| BKO:0000003 oxidoreductase activity | 2 (0) | 0 (0) | 2 (0) |
| BKO:0000002 hydroxylase activity | 2 (0) | 0 (0) | 2 (0) |
| BKO:0000018 protein hydroxylase activity | 2 (2) | 0 (0) | 2 (2) |
| BKO:0000198 catalytic activity, acting on a protein | 1870 (9) | 1097 (0) | 2967 (9) |
| BKO:0000028 catalytic activity, adding a chemical group on a protein | 1414 (34) | 515 (1) | 1929 (35) |
| BKO:0000265 protein acetyltransferase activity | 20 (20) | 1 (1) | 21 (21) |
| BKO:0000289 protein glycosyltransferase activity | 3 (3) | 0 (0) | 3 (3) |
| BKO:0000314 protein kinase activity | 1258 (1258) | 484 (484) | 1742 (1742) |
| BKO:0000327 protein methyltransferase activity | 1 (1) | 0 (0) | 1 (1) |
| BKO:0000339 protein palmitoyltransferase activity | 1 (1) | 0 (0) | 1 (1) |
| BKO:0000350 protein prenyltransferase activity | 1 (1) | 0 (0) | 1 (1) |
| BKO:0000374 ubiquitin-like protein transferase activity | 126 (0) | 30 (0) | 156 (0) |
| BKO:0000375 ubiquitin-protein transferase activity | 126 (126) | 30 (30) | 156 (156) |
| BKO:0000188 catalytic activity, removing a chemical group from a protein | 155 (4) | 59 (0) | 214 (4) |
| BKO:0000250 phosphoprotein phosphatase activity | 121 (121) | 57 (57) | 178 (178) |
| BKO:0000278 protein demethylase activity | 6 (6) | 0 (0) | 6 (6) |
| BKO:0000303 protein deacetylase activity | 11 (11) | 1 (1) | 12 (12) |
| BKO:0000362 ubiquitinyl hydrolase activity | 14 (0) | 2 (0) | 16 (0) |
| BKO:0000361 protein ubiquitinyl hydrolase activity | 14 (14) | 2 (2) | 16 (16) |
| BKO:0000206 catalytic activity, activating a protein | 604 (604) | 833 (833) | 1437 (1437) |
| BKO:0000636 catalytic activity, deactivating a protein | 75 (75) | 115 (115) | 190 (190) |
| BKO:0000185 hydrolase activity | 145 (0) | 59 (0) | 204 (0) |
| BKO:0000264 hydrolase activity, acting on ester bonds | 121 (0) | 57 (0) | 178 (0) |
| BKO:0000262 phosphoric ester hydrolase activity | 121 (0) | 57 (0) | 178 (0) |
| BKO:0000251 phosphatase activity | 121 (0) | 57 (0) | 178 (0) |
| BKO:0000250 phosphoprotein phosphatase activity | 121 (121) | 57 (57) | 178 (178) |
| BKO:0000304 deacetylase activity | 11 (0) | 1 (0) | 12 (0) |

| | | | |
|---|---|---|---|
| BKO:0000303 protein deacetylase activity | 11 (11) | 1 (1) | 12 (12) |
| BKO:0000362 ubiquitinyl hydrolase activity | 14 (0) | 2 (0) | 16 (0) |
| BKO:0000361 protein ubiquitinyl hydrolase activity | 14 (14) | 2 (2) | 16 (16) |
| BKO:0000211 transferase activity | 1380 (0) | 514 (0) | 1894 (0) |
| BKO:0000277 transferase activity, transferring acyl groups | 21 (0) | 1 (0) | 22 (0) |
| BKO:0000275 transferase activity, transferring acyl groups other than amino-acyl groups | 21 (0) | 1 (0) | 22 (0) |
| BKO:0000266 acetyltransferase activity | 20 (0) | 1 (0) | 21 (0) |
| BKO:0000265 protein acetyltransferase activity | 20 (20) | 1 (1) | 21 (21) |
| BKO:0000340 palmitoyltransferase activity | 1 (0) | 0 (0) | 1 (0) |
| BKO:0000339 protein palmitoyltransferase activity | 1 (1) | 0 (0) | 1 (1) |
| BKO:0000290 transferase activity, transferring glycosyl groups | 3 (0) | 0 (0) | 3 (0) |
| BKO:0000289 protein glycosyltransferase activity | 3 (3) | 0 (0) | 3 (3) |
| BKO:0000324 transferase activity, transferring phosphorus-containing groups | 1258 (0) | 484 (0) | 1742 (0) |
| BKO:0000315 kinase activity | 1258 (0) | 484 (0) | 1742 (0) |
| BKO:0000314 protein kinase activity | 1258 (1258) | 484 (484) | 1742 (1742) |
| BKO:0000337 transferase activity, transferring one-carbon groups | 1 (0) | 0 (0) | 1 (0) |
| BKO:0000328 methyltransferase activity | 1 (0) | 0 (0) | 1 (0) |
| BKO:0000327 protein methyltransferase activity | 1 (1) | 0 (0) | 1 (1) |
| BKO:0000351 transferase activity, transferring prenyl groups | 1 (0) | 0 (0) | 1 (0) |
| BKO:0000350 protein prenyltransferase activity | 1 (1) | 0 (0) | 1 (1) |
| BKO:0000374 ubiquitin-like protein transferase activity | 126 (0) | 30 (0) | 156 (0) |
| BKO:0000375 ubiquitin-protein transferase activity | 126 (126) | 30 (30) | 156 (156) |
| BKO:0000212 metabolic catalytic activity | 275 (275) | 802 (802) | 1077 (1077) |
| BKO:0000279 demethylase activity | 6 (0) | 0 (0) | 6 (0) |
| BKO:0000278 protein demethylase activity | 6 (6) | 0 (0) | 6 (6) |
| BKO:0000399 inhibitory activity | 1002 (1002) | 168 (168) | 1170 (1170) |
| BKO:0000077 transporter activity | 430 (430) | 151 (151) | 581 (581) |
| BKO:0000220 binding | 1590 (113) | 527 (25) | 2117 (138) |
| BKO:0000232 protein binding | 1477 (1477) | 502 (502) | 1979 (1979) |
| BKO:0000088 process | 9863 (0) | 3852 (0) | 13715 (0) |
| BKO:0000086 biochemical or transport reaction | 5117 (0) | 2978 (0) | 8095 (0) |
| BKO:0000085 transport reaction | 430 (0) | 151 (0) | 581 (0) |
| BKO:0000563 co-transport reaction | 4 (0) | 4 (0) | 8 (0) |
| BKO:0000577 symporter-mediated transport | 0 (0) | 4 (4) | 4 (4) |
| BKO:0000580 antiporter-mediated transport | 4 (4) | 0 (0) | 4 (4) |
| BKO:0000565 passive transport | 426 (426) | 151 (151) | 577 (577) |
| BKO:0000574 active transport | 1 (1) | 0 (0) | 1 (1) |
| BKO:0000103 biochemical reaction | 3719 (0) | 2630 (0) | 6349 (0) |

| | | | |
|---|---|---|---|
| BKO:0000101 conversion | 1409 (0) | 1015 (0) | 2424 (0) |
| BKO:0000099 removal of a chemical group | 179 (0) | 57 (0) | 236 (0) |
| BKO:0000098 deacetylation | 13 (0) | 1 (0) | 14 (0) |
| BKO:0000118 protein deacetylation | 13 (13) | 1 (1) | 14 (14) |
| BKO:0000112 deprenylation | 3 (0) | 0 (0) | 3 (0) |
| BKO:0000156 protein deprenylation | 3 (3) | 0 (0) | 3 (3) |
| BKO:0000116 deubiquitination | 20 (0) | 3 (0) | 23 (0) |
| BKO:0000168 protein deubiquitination | 20 (20) | 3 (3) | 23 (23) |
| BKO:0000119 removal of a chemical group from a protein | 179 (12) | 57 (0) | 236 (12) |
| BKO:0000118 protein deacetylation | 13 (13) | 1 (1) | 14 (14) |
| BKO:0000156 protein deprenylation | 3 (3) | 0 (0) | 3 (3) |
| BKO:0000168 protein deubiquitination | 20 (20) | 3 (3) | 23 (23) |
| BKO:0000409 protein demethylation | 6 (6) | 0 (0) | 6 (6) |
| BKO:0000414 protein dephosphorylation | 127 (127) | 54 (54) | 181 (181) |
| BKO:0000394 demethylation | 6 (0) | 0 (0) | 6 (0) |
| BKO:0000409 protein demethylation | 6 (6) | 0 (0) | 6 (6) |
| BKO:0000415 dephosphorylation | 127 (0) | 54 (0) | 181 (0) |
| BKO:0000414 protein dephosphorylation | 127 (127) | 54 (54) | 181 (181) |
| BKO:0000227 deactivation | 66 (66) | 108 (108) | 174 (174) |
| BKO:0000385 addition of a chemical group | 1039 (0) | 440 (0) | 1479 (0) |
| BKO:0000384 addition of a chemical group on a protein | 1039 (61) | 440 (1) | 1479 (62) |
| BKO:0000423 protein prenylation | 3 (3) | 0 (0) | 3 (3) |
| BKO:0000430 protein acetylation | 19 (19) | 1 (1) | 20 (20) |
| BKO:0000443 protein glycosylation | 5 (5) | 0 (0) | 5 (5) |
| BKO:0000450 protein hydroxylation | 2 (2) | 0 (0) | 2 (2) |
| BKO:0000460 protein methylation | 1 (1) | 0 (0) | 1 (1) |
| BKO:0000470 protein myristoylation | 1 (1) | 0 (0) | 1 (1) |
| BKO:0000480 protein palmitoylation | 2 (2) | 0 (0) | 2 (2) |
| BKO:0000490 protein phosphorylation | 835 (835) | 406 (406) | 1241 (1241) |
| BKO:0000520 protein ubiquitination | 122 (122) | 33 (33) | 155 (155) |
| BKO:0000396 prenylation | 3 (0) | 0 (0) | 3 (0) |
| BKO:0000423 protein prenylation | 3 (3) | 0 (0) | 3 (3) |
| BKO:0000431 acetylation | 19 (0) | 1 (0) | 20 (0) |
| BKO:0000430 protein acetylation | 19 (19) | 1 (1) | 20 (20) |
| BKO:0000441 glycosylation | 5 (0) | 0 (0) | 5 (0) |
| BKO:0000443 protein glycosylation | 5 (5) | 0 (0) | 5 (5) |
| BKO:0000451 hydroxylation | 2 (0) | 0 (0) | 2 (0) |
| BKO:0000450 protein hydroxylation | 2 (2) | 0 (0) | 2 (2) |
| BKO:0000461 methylation | 1 (0) | 0 (0) | 1 (0) |
| BKO:0000460 protein methylation | 1 (1) | 0 (0) | 1 (1) |
| BKO:0000471 myristoylation | 1 (0) | 0 (0) | 1 (0) |
| BKO:0000470 protein myristoylation | 1 (1) | 0 (0) | 1 (1) |
| BKO:0000481 palmitoylation | 2 (0) | 0 (0) | 2 (0) |
| BKO:0000480 protein palmitoylation | 2 (2) | 0 (0) | 2 (2) |
| BKO:0000491 phosphorylation | 835 (0) | 406 (0) | 1241 (0) |
| BKO:0000490 protein phosphorylation | 835 (835) | 406 (406) | 1241 (1241) |
| BKO:0000521 ubiquitination | 122 (0) | 33 (0) | 155 (0) |
| BKO:0000520 protein ubiquitination | 122 (122) | 33 (33) | 155 (155) |
| BKO:0000567 activation | 340 (340) | 761 (761) | 1101 (1101) |
| BKO:0000175 dissociation | 226 (50) | 190 (56) | 416 (106) |
| BKO:0000174 protein complex dissociation | 176 (166) | 134 (123) | 310 (289) |
| BKO:0000182 protein multimer dissociation | 10 (10) | 11 (11) | 21 (21) |
| BKO:0000225 non-covalent binding | 1641 (275) | 571 (172) | 2212 (447) |

| | | | |
|---|---|---|---|
| BKO:0000241 protein complex formation | 1366 (1315) | 399 (355) | 1765 (1670) |
| BKO:0000240 protein multimerisation | 51 (51) | 44 (44) | 95 (95) |
| BKO:0000530 degradation | 230 (230) | 63 (63) | 293 (293) |
| BKO:0000535 conformational transition | 6 (0) | 50 (0) | 56 (0) |
| BKO:0000582 channel closing | 0 (0) | 6 (6) | 6 (6) |
| BKO:0000585 channel opening | 6 (6) | 44 (44) | 50 (50) |
| BKO:0000589 metabolic reaction | 334 (0) | 869 (0) | 1203 (0) |
| BKO:0000588 irreversible metabolic reaction | 325 (325) | 868 (868) | 1193 (1193) |
| BKO:0000591 reversible metabolic reaction | 9 (9) | 1 (1) | 10 (10) |
| BKO:0000545 translocation reaction | 1085 (1085) | 337 (337) | 1422 (1422) |
| BKO:0000131 irreversible process | 8911 (8091) | 3548 (2257) | 12459 (10348) |
| BKO:0000530 degradation | 230 (230) | 63 (63) | 293 (293) |
| BKO:0000588 irreversible metabolic reaction | 325 (325) | 868 (868) | 1193 (1193) |
| BKO:0000615 omitted irreversible process | 295 (295) | 377 (377) | 672 (672) |
| BKO:0000216 uncertain process | 518 (0) | 48 (0) | 566 (0) |
| BKO:0000215 uncertain irreversible process | 518 (518) | 47 (47) | 565 (565) |
| BKO:0000219 uncertain reversible process | 518 (518) | 48 (48) | 566 (566) |
| BKO:0000249 molecular or genetic interaction | 1366 (0) | 399 (0) | 1765 (0) |
| BKO:0000244 molecular interaction | 1366 (0) | 399 (0) | 1765 (0) |
| BKO:0000241 protein complex formation | 1366 (1315) | 399 (355) | 1765 (1670) |
| BKO:0000240 protein multimerisation | 51 (51) | 44 (44) | 95 (95) |
| BKO:0000537 composite biochemical process | 1900 (0) | 188 (0) | 2088 (0) |
| BKO:0000536 transcription | 970 (970) | 138 (138) | 1108 (1108) |
| BKO:0000541 translation | 930 (930) | 50 (50) | 980 (980) |
| BKO:0000551 biological process | 504 (504) | 148 (148) | 652 (652) |
| BKO:0000556 omitted process | 295 (0) | 379 (0) | 674 (0) |
| BKO:0000615 omitted irreversible process | 295 (295) | 377 (377) | 672 (672) |
| BKO:0000617 omitted reversible process | 0 (0) | 2 (2) | 2 (2) |
| BKO:0000561 reversible process | 18 (9) | 5 (2) | 23 (11) |
| BKO:0000591 reversible metabolic reaction | 9 (9) | 1 (1) | 10 (10) |
| BKO:0000617 omitted reversible process | 0 (0) | 2 (2) | 2 (2) |
| **Total** | **109681 (31087)** | **49986 (13198)** | **159667 (44285)** |

Table S3: **List of the 30 template bricks representing generic concepts.** For each template brick, we also give the number of instances they match in the ACSN and PANTHER databases. The number in parantheses is th number of proper instances matched by the template brick, i.e. the number of instances that are matched by the template brick and by none of its descendants.

| **Template brick** | **ACSN** | **PANTHER** | **Overall** |
|---|---|---|---|
| BKO:0000130 irreversible process (PD main) | 8911 (2605) | 3548 (251) | 12459 (2856) |
| BKO:0000387 addition of a chemical group on a protein (PD narrow 1) | 791 (54) | 364 (1) | 1155 (55) |
| BKO:0000389 addition of a chemical group on a protein (PD narrow 2) | 230 (6) | 67 (0) | 297 (6) |
| BKO:0000391 addition of a chemical group on a protein (PD narrow 3) | 10 (0) | 2 (0) | 12 (0) |
| BKO:0000390 addition of a chemical group on a protein (PD narrow 4) | 9 (1) | 8 (0) | 17 (1) |
| BKO:0000126 removal of a chemical group from a protein (PD narrow 1) | 122 (11) | 51 (0) | 173 (11) |
| BKO:0000127 removal of a chemical group from a protein (PD narrow 2) | 56 (1) | 5 (0) | 61 (1) |
| BKO:0000129 removal of a chemical group from a protein (PD narrow 3) | 0 (0) | 1 (0) | 1 (0) |

| | | | |
|---|---|---|---|
| BKO:0000128 removal of a chemical group from a protein (PD narrow 4) | 1 (0) | 0 (0) | 1 (0) |
| BKO:0000087 modulatory activity (PD narrow 1) | 5928 (60) | 2584 (0) | 8512 (60) |
| BKO:0000009 stimulatory activity (PD narrow 1) | 4965 (267) | 2429 (18) | 7394 (285) |
| BKO:0000006 catalytic activity (PD narrow 1) | 4681 (2544) | 2410 (512) | 7091 (3056) |
| BKO:0000027 catalytic activity, acting on a protein (PD narrow 1) | 1495 (5) | 999 (0) | 2494 (5) |
| BKO:0000023 catalytic activity, adding a chemical group on a protein (PD narrow 1) | 1138 (29) | 438 (1) | 1576 (30) |
| BKO:0000199 catalytic activity, removing a chemical group from a protein (PD narrow 1) | 110 (4) | 50 (0) | 160 (4) |
| BKO:0000029 catalytic activity, acting on a protein (PD narrow 2) | 315 (4) | 82 (0) | 397 (4) |
| BKO:0000024 catalytic activity, adding a chemical group on a protein (PD narrow 2) | 256 (1) | 68 (0) | 324 (1) |
| BKO:0000200 catalytic activity, removing a chemical group from a protein (PD narrow 2) | 45 (0) | 8 (0) | 53 (0) |
| BKO:0000205 catalytic activity, acting on a protein (PD narrow 3) | 10 (0) | 3 (0) | 13 (0) |
| BKO:0000026 catalytic activity, adding a chemical group on a protein (PD narrow 3) | 10 (0) | 2 (0) | 12 (0) |
| BKO:0000202 catalytic activity, removing a chemical group from a protein (PD narrow 3) | 0 (0) | 1 (0) | 1 (0) |
| BKO:0000204 catalytic activity, acting on a protein (PD narrow 4) | 11 (0) | 9 (0) | 20 (0) |
| BKO:0000025 catalytic activity, adding a chemical group on a protein (PD narrow 4) | 11 (2) | 8 (0) | 19 (2) |
| BKO:0000201 catalytic activity, removing a chemical group from a protein (PD narrow 4) | 0 (0) | 0 (0) | 0 (0) |
| BKO:0000081 transport reaction (PD narrow 1) | 420 (0) | 109 (0) | 529 (0) |
| BKO:0000082 transport reaction (PD narrow 2) | 330 (0) | 95 (0) | 425 (0) |
| BKO:0000097 modulatory activity (PD narrow 2) | 11 (0) | 6 (0) | 17 (0) |
| BKO:0000017 stimulatory activity (PD narrow 2) | 11 (0) | 6 (0) | 17 (0) |
| BKO:0000007 catalytic activity (PD narrow 2) | 11 (3) | 6 (5) | 17 (8) |
| BKO:0000562 reversible process (PD main) | 18 (9) | 5 (2) | 23 (11) |
| **Total** | **29906 (5606)** | **13364 (790)** | **43270 (6396)** |

Table S4: **Classification of all instances of the ACSN and PANTHER databases that were only matched by template bricks representing generic concepts.** All bricks that were matched by one of the 30 template bricks listed below and representing generic concepts (e.g. an irreversible process, a catalytic activity) without being matched by template bricks representing more specific concepts (e.g. a protein phosphorylation, a protein kinase activity) were classified in one of the following categories and subcategories. **Misrepresentation**: the process or activity is badly represented; *Same reactant and product*: a process consumes a reactant and produces a product that are exactly the same entity pools; *Phenotype as modulator*: a phenotype modulates a process (invalid SBGN); *Phenotype as reactant or product*: a phenotype is a reactant or a product of a process (invalid SBGN); *Incomplete process*: the process is missing some reactant or product; Association: the process is an incomplete association; Dissociation: the process is an incomplete dissociation; *False multimer*: a multimer is represented using two subunits on top of each other and not a multimer glyph; *Transcription no gene ui*: a transcription is represented without using a proper unit of information for the gene; *Translation no ui*: a translation is represented without using a proper unit of information for the mRNA; *Other*: none of the above. **Nature of process implicit in label**: the nature of the process can solely be identified by analyzing the labels of the participants of the process (e.g. the truncation of a protein, where a protein is consumed to give two or more proteins with new names); *Truncation*: the process is a truncation; *Translocation*: the process is a translocation. *Other*: none of the above. **Process in the ontology**: this category implies only modulatory activities whose modulated process is an instance of a template brick that belongs to the ontology (e.g. stimulation of dissociation). *Activation*: the process is an activation; *Translocation*: the process is a translocation; *Complexation*: the process is a complexation; *Degradation*: the process is a degradation; *Dissociation*: the process is a dissociation; *Phosphorylation*: the process is a phosphorylation; *Transcription*: the process is a transcription; *Translation*: the process is a translation; *Ubiquitination*: the process is an ubiquitination; *Association*: the process is an association; *Multimer dissociation*: the process is a multimer dissociation; *Multimerisation*: the process is a multimerisation; *Transporter activity*: the process is a translocation; *Channel opening*: the process is a channel opening. *Other*: none of the above. **Mix of two processes of unknown nature**: two processes are represented using only one SBGN process (e.g. a dissociation and a dephosphorylation); *Association and deactivation*: the process is an association and a deactivation; *Association and multimerisation*: the process is an association and a multimerisation; *Dissociation and activation*: the process is a dissociation and an activation; *Dissociation and deubiquitination*: the process is a dissociation and a deubiquitination; *Dissociation and multimer dissociation*: the process is a dissociation and a multimer dissociation; *Dissociation and phosphorylation*: the process is a dissociation and a phosphorylation; *Dissociation and ubiquitination*: the process is a dissociation and an ubiquitination; *Association and phosphorylation*: the process is an association and a phosphorylation; *Association and activation*: the process is an association and a activation; *Dissociation and acetylation*: the process is a dissociation and an acetylation; *Dissociation and dephosphorylation*: the process is a dissociation and a dephosphorylation; *Multimer dissociation and deactivation*: the process is a multimer dissociation and a deactivation; *Multimerisation and activation*: the process is a multimerisation and an activation; *Multimerisation and ubiquitination*: the process is a multimerisation and an ubiquitination. *Multimerisation and deacetylation*: the process is a multimerisation and a deacetylation; *Multimerisation and phosphorylation*: the process is a multimerisation and a phosphorylation; **Process of unknown nature**: we could not identify the nature of the represented process or the activity. **Process not in the ontology**: the process represented in the brick is not an instance of any template brick of the ontology; *Non standard value in state variable*: the process sets a value to a state variable that is not defined by the controlled vocabulary of the SBGN standard; Asterisk: the value is an asterisk ("*"). This value is used in CellDesigner to mean "don't care". It does not belong to the controlled vocabulary of SBGN, but its use in state variables is valid SBGN. However processes using this value cannot be linked to any specific concept of SBO or GO. Question mark: the value is a question mark ("?"). Same as for "*", but with meaning "unknown". Other: none of the above; *Process involving an asRNA*: the process consumes or produces an asRNA; *Methylation of a naf of unspecified nature*: the process is a methylation of a nuclear acid feature with no conceptual type defined (e.g. gene or mRNA); *Gtp exchange*: the process is a GTP/GDP exchange process; *Production*: the process consumes an empty set.

| Template brick | Category | Subcategories | | ACSN | PANTHER | Overall |
|---|---|---|---|---|---|---|
| catalytic activity, removing a chemical group from a protein (PD narrow 1) | misrepresentation | other | | 1 | 0 | 1 |
| | process not in the ontology | non standard value in state variable | asterisk | 2 | 0 | 2 |
| | | | question mark | 1 | 0 | 1 |
| catalytic activity, adding a chemical group on a protein (PD narrow 1) | misrepresentation | other | | 1 | 0 | 1 |
| | nature of process implicit in label | truncation | | 3 | 0 | 3 |
| | process not in the ontology | non standard value in state variable | asterisk | 3 | 0 | 3 |
| | | | question mark | 22 | 1 | 23 |
| addition of a chemical group on a protein (PD narrow 1) | nature of process implicit in label | truncation | | 2 | 0 | 2 |
| | process not in the ontology | non standard value in state variable | asterisk | 6 | 0 | 6 |
| | | | other | 4 | 0 | 4 |
| | | | question mark | 42 | 1 | 43 |
| stimulatory activity (PD narrow 1) | misrepresentation | phenotype as modulator | | 4 | 0 | 4 |
| | | phenotype as reactant or product | | 7 | 0 | 7 |
| | | other | | 0 | 1 | 1 |
| | | same reactant and product | | 30 | 0 | 30 |
| | nature of process implicit in label | other | | 14 | 0 | 14 |
| | | truncation | | 1 | 0 | 1 |
| | process in the ontology | activation | | 0 | 12 | 12 |
| | | complexation | | 36 | 0 | 36 |
| | | degradation | | 2 | 0 | 2 |
| | | dissociation | | 10 | 0 | 10 |
| | | phosphorylation | | 23 | 1 | 24 |
| | | transcription | | 125 | 4 | 129 |
| | | translation | | 6 | 0 | 6 |
| | | ubiquitination | | 3 | 0 | 3 |
| | process not in the ontology | non standard value in state variable | other | 1 | 0 | 1 |
| | | | question mark | 4 | 0 | 4 |
| | | process involving an asrna | | 1 | 0 | 1 |
| addition of a chemical group on a protein | process not in the ontology | non standard value in state variable | asterisk | 3 | 0 | 3 |
| | | | question mark | 3 | 0 | 3 |
| (PD narrow 2) | misrepresentation | incomplete process | association | 27 | 8 | 35 |
| | | | dissociation | 24 | 2 | 26 |
| | | phenotype as reactant or product | | 1120 | 7 | 1127 |
| | | false multimer | | 7 | 0 | 7 |

| | | | | | | |
|---|---|---|---|---|---|---|
| | | other | | 12 | 15 | 27 |
| | | same reactant and product | | 141 | 19 | 160 |
| | | transcription no gene ui | | 3 | 0 | 3 |
| | | translation no ui | | 2 | 0 | 2 |
| | mix of two processes of different nature | association and deactivation | | 0 | 1 | 1 |
| | | association and multimerisation | | 16 | 2 | 18 |
| | | dissociation and activation | | 0 | 2 | 2 |
| | | dissociation and deubiquitination | | 0 | 1 | 1 |
| | | dissociation and multimer dissociation | | 0 | 2 | 2 |
| | | dissociation and phosphorylation | | 0 | 1 | 1 |
| | | dissociation and ubiquitination | | 0 | 1 | 1 |
| | nature of process implicit in label | other | | 1001 | 110 | 1111 |
| | | translocation | | 0 | 2 | 2 |
| | nature of process implicit in label | truncation | | 122 | 62 | 184 |
| | process not in the ontology | methylation of a naf of unspecified nature | | 2 | 0 | 2 |
| | | gtp exchange | | 25 | 10 | 35 |
| | | production | | 0 | 1 | 1 |
| | | non standard value in state variable | asterisk | 1 | 0 | 1 |
| | | | other | 19 | 0 | 19 |
| | | | question mark | 20 | 0 | 20 |
| | | process involving an as-rna | | 56 | 0 | 56 |
| | process of unknown nature | | | 7 | 5 | 12 |
| modulatory activity (PD narrow 1) | misrepresentation | phenotype as modulator | | 4 | 0 | 4 |
| | | phenotype as reactant or product | | 2 | 0 | 2 |
| | mix of two processes of different nature | association and phosphorylation | | 1 | 0 | 1 |
| | nature of process implicit in label | other | | 28 | 0 | 28 |
| | | truncation | | 1 | 0 | 1 |

| | | | | | | |
|---|---|---|---|---|---|---|
| | process in the ontology | complexation | | 3 | 0 | 3 |
| | | degradation | | 2 | 0 | 2 |
| | | other | | 14 | 0 | 14 |
| | | transcription | | 5 | 0 | 5 |
| catalytic activity (PD narrow 1) | misrepresentation | incomplete process | association | 11 | 0 | 11 |
| | | | dissociation | 11 | 0 | 11 |
| | | phenotype as modulator | | 8 | 0 | 8 |
| | | phenotype as reactant or product | | 74 | 0 | 74 |
| | | false multimer | | 10 | 0 | 10 |
| | | other | | 4 | 4 | 8 |
| | | same reactant and product | | 74 | 6 | 80 |
| | mix of two processes of different nature | association and activation | | 5 | 4 | 9 |
| | | association and deactivation | | 5 | 0 | 5 |
| | | association and multimerisation | | 4 | 0 | 4 |
| | | association and phosphorylation | | 4 | 0 | 4 |
| | | dissociation and acetylation | | 1 | 0 | 1 |
| | | dissociation and activation | | 0 | 1 | 1 |
| | | dissociation and dephosphorylation | | 2 | 0 | 2 |
| | | multimer dissociation and deactivation | | 0 | 4 | 4 |
| | | multimerisation and activation | | 8 | 2 | 10 |
| | | multimerisation and ubiquitination | | 1 | 0 | 1 |
| | | multimersation and deacetylation | | 1 | 0 | 1 |
| | | multimerisation and phosphorylation | | 0 | 4 | 4 |
| | nature of process implicit in label | other | | 280 | 42 | 322 |
| | | translocation | | 0 | 5 | 5 |
| | | truncation | | 151 | 70 | 221 |
| | | association | | 171 | 50 | 221 |
| | | degradation | | 62 | 23 | 85 |

process in the ontology

| | | | | | | |
|---|---|---|---|---|---|---|
| | | dissociation | | 28 | 71 | 99 |
| | | multimer dissociation | | 2 | 0 | 2 |
| | | multimerisation | | 14 | 9 | 23 |
| | | transcription | | 1100 | 173 | 1273 |
| | | translation | | 75 | 0 | 75 |
| | | transporter activity | | 324 | 16 | 340 |
| | process not in the ontology | methylation of a naf of unspecified nature | | 2 | 0 | 2 |
| | | gtp exchange | | 55 | 25 | 80 |
| | | non standard value in state variable | other | 7 | 0 | 7 |
| | | | question mark | 1 | 0 | 1 |
| | | process involving an as-rna | | 45 | 0 | 45 |
| | | production | | 2 | 2 | 4 |
| | process of unknown nature | | | 2 | 1 | 3 |
| catalytic activity, adding a chemical group on a protein (PD narrow 2) | process not in the ontology | non standard value in state variable | question mark | 1 | 0 | 1 |
| catalytic activity, acting on a protein (PD narrow 1) | misrepresentation | other | | 1 | 0 | 1 |
| | process in the ontology | channel opening | | 2 | 0 | 2 |
| | process not in the ontology | non standard value in state variable | other | 2 | 0 | 2 |
| catalytic activity (PD narrow 2) | misrepresentation | phenotype as modulator | | 2 | 0 | 2 |
| | | other | | 0 | 4 | 4 |
| | nature of process implicit in label | other | | 1 | 1 | 2 |
| removal of a chemical group from a protein (PD narrow 2) | process not in the ontology | non standard value in state variable | question mark | 1 | 0 | 1 |
| removal of a chemical group from a protein (PD narrow 1) | process not in the ontology | non standard value in state variable | asterisk | 3 | 0 | 3 |
| | | | other | 2 | 0 | 2 |
| | | | question mark | 6 | 0 | 6 |
| catalytic activity, acting on a protein (PD narrow 2) | misrepresentation | same reactant and product | | 4 | 0 | 4 |
| addition of a chemical group on a protein (PD narrow 4) | process not in the ontology | non standard value in state variable | question mark | 1 | 0 | 1 |

| | | | | | | |
|---|---|---|---|---|---|---|
| catalytic activity, adding a chemical group on a protein (PD narrow 4) | process not in the ontology | non standard value in state variable | question mark | 2 | 0 | 2 |
| **Total** | | | | **5606** | **790** | **6396** |

Table S5: **Classification of all instances of the ACSN and PANTHER databases that were only matched by template bricks representing generic concepts.** All bricks that were matched by one of 30 template bricks representing generic concepts (e.g. an irreversible process, a catalytic activity) without being matched by template bricks representing more specific concepts (e.g. a protein phosphorylation, a protein kinase activity) were classified in one of the following categories and subcategories. **Misrepresentation**: the process or activity is badly represented; *Same reactant and product*: a process consumes a reactant and produces a product that are exactly the same entity pools; *Phenotype as modulator*: a phenotype modulates a process (invalid SBGN); *Phenotype as reactant or product*: a phenotype is a reactant or a product of a process (invalid SBGN); *Incomplete process*: the process is missing some reactant or product; Association: the process is an incomplete association; Dissociation: the process is an incomplete dissociation; *False multimer*: a multimer is represented using two subunits on top of each other and not a multimer glyph; *Transcription no gene ui*: a transcription is represented without using a proper unit of information for the gene; *Translation no ui*: a translation is represented without using a proper unit of information for the mRNA; *Other*: none of the above. **Nature of process implicit in label**: the nature of the process can solely be identified by analyzing the labels of the participants of the process (e.g. the truncation of a protein, where a protein is consumed to give two or more proteins with new names); *Truncation*: the process is a truncation; *Translocation*: the process is a translocation. *Other*: none of the above. **Process of unknown nature**: we could not identify the nature of the represented process or the activity. **Process in the ontology**: this category implies only modulatory activities whose modulated process is an instance of a template brick that belongs to the ontology (e.g. stimulation of dissociation). *Activation*: the process is an activation; *Translocation*: the process is a translocation; *Complexation*: the process is a complexation; *Degradation*: the process is a degradation; *Dissociation*: the process is a dissociation; *Phosphorylation*: the process is a phosphorylation; *Transcription*: the process is a transcription; *Translation*: the process is a translation; *Ubiquitination*: the process is an ubiquitination; *Association*: the process is an association; *Multimer dissociation*: the process is a multimer dissociation; *Multimerisation*: the process is a multimerisation; *Transporter activity*: the process is a translocation; *Channel opening*: the process is a channel opening. *Other*: none of the above. **Mix of two processes of unknown nature**: two processes are represented using only one SBGN process (e.g. a dissociation and a dephosphorylation); *Association and deactivation*: the process is an association and a deactivation; *Association and multimerisation*: the process is an association and a multimerisation; *Dissociation and activation*: the process is a dissociation and an activation; *Dissociation and deubiquitination*: the process is a dissociation and a deubiquitination; *Dissociation and multimer dissociation*: the process is a dissociation and a multimer dissociation; *Dissociation and phosphorylation*: the process is a dissociation and a phosphorylation; *Dissociation and ubiquitination*: the process is a dissociation and an ubiquitination; *Association and phosphorylation*: the process is an association and a phosphorylation; *Association and activation*: the process is an association and a activation; *Dissociation and acetylation*: the process is a dissociation and an acetylation; *Dissociation and dephosphorylation*: the process is a dissociation and a dephosphorylation; *Multimer dissociation and deactivation*: the process is a multimer dissociation and a deactivation; *Multimerisation and activation*: the process is a multimerisation and an activation; *Multimerisation and ubiquitination*: the process is a multimerisation and an ubiquitination. *Multimerisation and deacetylation*: the process is a multimerisation and a deacetylation; *Multimerisation and phosphorylation*: the process is a multimerisation and a phosphorylation; **Process not in the ontology**: the process represented in the brick is not an instance of any template brick of the ontology; *Non standard value in state variable*: the process sets a value to a state variable that is not defined by the controlled vocabulary of the SBGN standard; Asterisk: the value is an asterisk ("*"). This value is used in CellDesigner to mean "don't care". It does not belong to the controlled vocabulary of SBGN, but its use in state variables is valid SBGN. However processes using this value cannot be linked to any specific concept of SBO or GO. Question mark: the value is a question mark ("?"). Same as for "*", but with meaning "unknown". Other: none of the above; *Process involving an asRNA*: the process consumes or produces an asRNA; *Methylation of a naf of unspecified nature*: the process is a methylation of a nuclear acid feature with no conceptual type defined (e.g. gene or mRNA); *Gtp exchange*: the process is a GTP/GDP exchange process; *Production*: the process consumes an empty set.

| **Category** | **Subcategories** | | **ACSN** | **PANTHER** | **Overall** |
|---|---|---|---|---|---|
| misrepresentation | same reactant and product | | 252 | 25 | 277 |
| | phenotype as modulator | | 18 | 0 | 18 |
| | phenotype as reactant or product | | 1203 | 7 | 1210 |
| | incomplete process | association | 38 | 8 | 46 |
| | | dissociation | 35 | 2 | 37 |
| | false multimer | | 17 | 0 | 17 |
| | transcription no gene ui | | 3 | 0 | 3 |
| | translation no ui | | 2 | 0 | 2 |
| | other | | 19 | 24 | 43 |
| nature of process implicit in label | truncation | | 280 | 132 | 412 |
| | translocation | | 0 | 7 | 7 |

| | | | | | |
|---|---|---|---|---|---|
| | other | | 1328 | 154 | 1482 |
| process of unknown nature | | | 9 | 6 | 15 |
| process in the ontology | activation | | 0 | 13 | 13 |
| | translocation | | 2 | 0 | 2 |
| | complexation | | 39 | 0 | 39 |
| | degradation | | 66 | 23 | 89 |
| | dissociation | | 38 | 71 | 109 |
| | phosphorylation | | 23 | 1 | 24 |
| | transcription | | 1230 | 177 | 1407 |
| | translation | | 81 | 0 | 81 |
| | ubiquitination | | 3 | 0 | 3 |
| | association | | 171 | 50 | 221 |
| | multimer dissociation | | 2 | 0 | 2 |
| | multimerisation | | 14 | 9 | 23 |
| | transporter activity | | 324 | 16 | 340 |
| | channel opening | | 2 | 0 | 2 |
| | other | | 14 | 0 | 14 |
| mix of two processes of different nature | association and deactivation | | 5 | 1 | 6 |
| | association and multimerisation | | 20 | 2 | 22 |
| | dissociation and activation | | 0 | 3 | 3 |
| | dissociation and deubiquitination | | 0 | 1 | 1 |
| | dissociation and multimer dissociation | | 0 | 2 | 2 |
| | dissociation and phosphorylation | | 0 | 1 | 1 |
| | dissociation and ubiquitination | | 0 | 1 | 1 |
| | association and phosphorylation | | 5 | 0 | 5 |
| | association and activation | | 5 | 4 | 9 |
| | dissociation and acetylation | | 1 | 0 | 1 |
| | dissociation and dephosphorylation | | 2 | 0 | 2 |
| | multimer dissociation and deactivation | | 0 | 4 | 4 |
| | multimerisation and activation | | 8 | 2 | 10 |
| | multimerisation and ubiquitination | | 1 | 0 | 1 |
| | multimersation and deacetylation | | 1 | 0 | 1 |
| | multimerisation and phosphorylation | | 0 | 4 | 4 |
| process not in the ontology | non standard value in state variable | asterisk | 18 | 0 | 18 |
| | | question mark | 104 | 2 | 106 |
| | | other | 35 | 0 | 35 |
| | process involving an asrna | | 102 | 0 | 102 |
| | methylation of a naf of unspecified nature | | 4 | 0 | 4 |
| | gtp exchange | | 80 | 35 | 115 |
| | production | | 2 | 3 | 5 |
| **Total** | | | **5606** | **790** | **6396** |

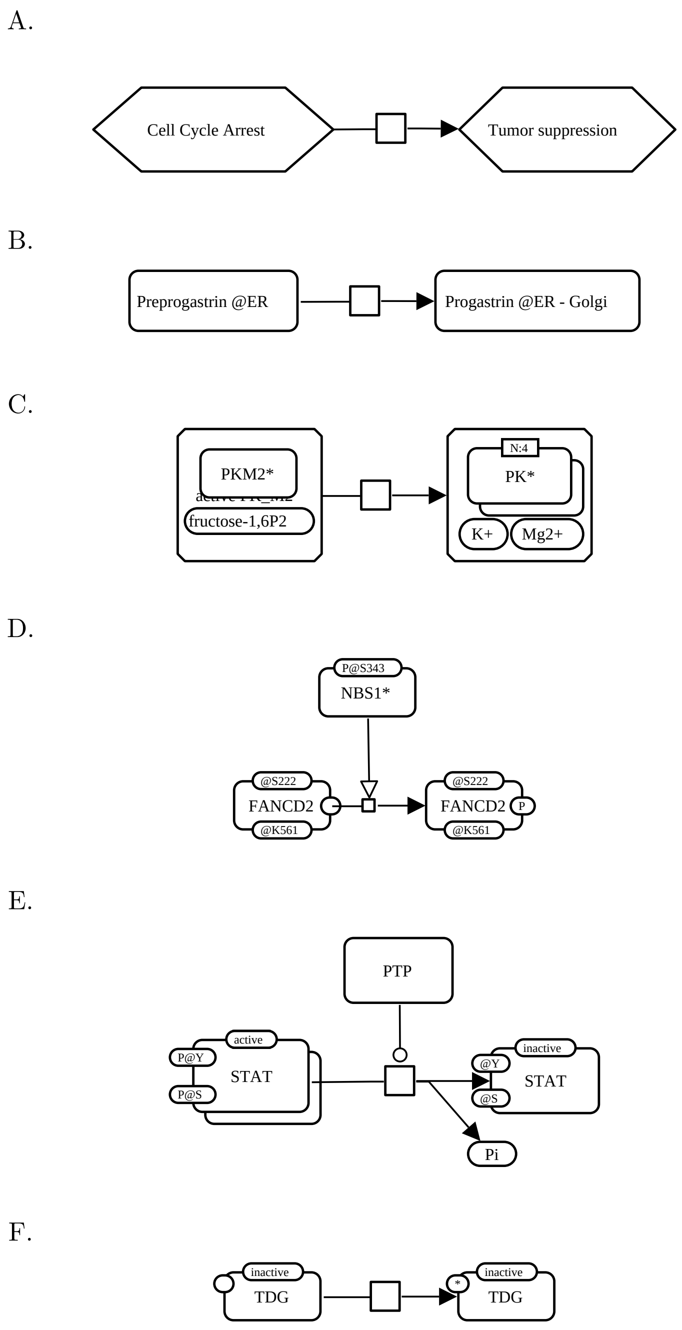


Figure S3: **Example of instance bricks for each main category.** A. Misrepresentation: the instance represents the consumption and production of phenotypes, which is invalid SBGN. It was retrieved from the "p53 pathway" map of the PANTHER database. B. Nature of process implicit in labels: the instance potentially represents the translocation of preprogastrin from the endoplasmic reticulum to the Golgi apparatus. It was retrieved from the "CCKR signaling map" of the PANTHER database. C. Process of unknown nature. The instance was retrieved from the "RCD master" map of the ACSN. D. Process in the ontology: the instance represents the stimulation of the phophorylation of FANCD2 bu NBS1. It was retrieved from the "Cell Cycle DNA repair master" map of the ACSN. E. Mix of two process of different nature: the instance represents the dephosphorylation and deactivation of STAT, catalyzed bu PTP. It was retrieved from the "JAK STAT signaling pathway" map of the ACSN. F. Process not in the ontology: the instance represents the modification of TDG to a state that is not indicated ("*" state variable value). It was retrieved from the "Cell Cycle DNA repair master" map of the ACSN.
